# Explainable Generative AI Uncovers a Molecular Continuum in Medulloblastoma with Implications for Rare Cancer Subtyping and Treatment Equity

**DOI:** 10.1101/2024.12.30.630738

**Authors:** Guillermo Prol-Castelo, Alejandro Tejada-Lapuerta, Beatriz Urda-García, Iker Núñez-Carpintero, Erica Garcia-Verellen, Arnau Montagud, Alfonso Valencia, Davide Cirillo

## Abstract

Medulloblastoma is a childhood brain tumor traditionally classified into four molecular subgroups. Recent evidence suggests that Groups 3 and 4 represent a biological continuum rather than distinct entities, a paradigm shift with significant implications for understanding disease biology and treatment strategies. Nevertheless, assessing this hypothesis is challenging mainly due to data scarcity. In this study, we analyze the largest available transcriptomics dataset to provide compelling evidence for the existence of an intermediate subgroup between Groups 3 and 4, characterized by distinct molecular features. To overcome limitations posed by data scarcity, we employ synthetic data generation using a Variational Autoencoder and apply explainability techniques to identify key relationships between gene expression and disease subgroups. Furthermore, by incorporating Machine Learning Fairness approaches, we demonstrate that overlooking this intermediate subgroup can result in treatment disparities. Our findings are further supported by both existing and newly proposed studies using diverse datasets and methodologies, including graphbased analyses and multi-scale simulations, underscoring the robustness and reproducibility of our results. This study demonstrates the potential of synthetic data generation to refine rare disease subtyping and advance our understanding of the underlying biological mechanisms.

Keywords: Medulloblastoma, pediatric cancer, representation learning, autoencoder, synthetic data

## 1. Introduction

Medulloblastoma (MB), the most common childhood brain tumor, is a highly heterogeneous disease with distinct subgroups characterized by differences in demographics, gene mutation patterns, and pathway associations (1, 2, 3, 4, 5). In 2012 (6), a consensus classification divided MB into four molecularly defined subgroups: Wingless (WNT), Sonic Hedgehog (SHH), Group 3 (G3), and Group 4 (G4). The consensus was updated in 2016 (7) and then again in 2021 (8). The latest consensus reflects the subgroup-defining mutations of WNT and SHH and the high variability observed within G3 and G4 in several studies (9, 10, 11, 12), dividing MB into WNT, SHH and non-WNT/non-SHH subgroups. Furthermore, studies have also suggested the existence of additional subgroups (3, 9), and specifically a subgroup with molecular features intermediate to those of G3 and G4 (13). In particular, our previous study using multi-omics data and a multilayer network methodology identified a putative G3-G4 subgroup (14), termed G3-G4, providing initial evidence for this intermediate classification.

Indeed, identifying this putative subgroup between G3 and G4 has been the subject of increasing interest given the differences in prognosis and the potential to provide personalized treatments. In (10), the authors found an overlap between G3 and G4 when using microarray and methylation data independently. This overlap was less clear when both datasets were combined, highlighting complexities that warrant further investigation. More recent studies do point out the possible existence of an intermediate group with characteristics between G3 and G4 when considering proteomic data (15), risk stratification (9), and multi-omics data (16). In particular, our previous study (14), using a multi-layer network approach, identified patients with intermediate characteristics between G3 and G4. Recently, a scoring system has been proposed to analyze the G3 and G4 continuum (17), with the authors themselves noting limitations due to data scarcity.

MB occurs approximately in 5 cases per 1 million in the pediatric population (18), with varying incidence among subgroups based on age. Scarcity of data is a common challenge in rare disease research, which is further complicated when cohorts are disaggregated by specific attributes (e.g., demographics, clinical or molecular characteristics, etc.), which can render statistical methods less effective (19). To mitigate these limitations, this study employs the largest publicly available MB omics dataset, comprising 763 samples (10) (see Table 1).

**Table 1.**
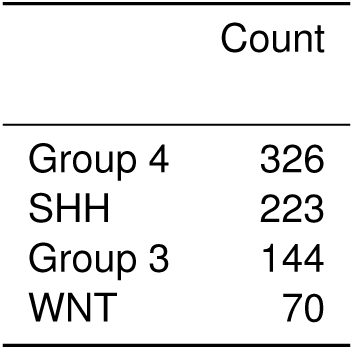
Patient distribution across Medulloblastoma subgroups. This table presents the number of patients in each medulloblastoma group, based on the data from 11. The table also shows the subgroup imbalance within the dataset.

In recent years, synthetic data generation (SDG) has emerged as a promising strategy to address the challenges posed by data scarcity by creating artificial data that mimics real-world data for analysis or modeling. This approach has proven particularly valuable in the biomedical domain, where data is often limited and subject to strict privacy constraints (20). The Variational Autoencoder (VAE) (21) is an unsupervised method that leverages both qualities of representation learning and generative models, resulting in a widespread use in cases of data scarcity and imbalance (22, 23). More specifically, VAEs have been used widely to learn embedded representations of transcriptomic data types, including bulk (24, 25), single-cell (26, 27), and spatial (28) transcriptomics, with several applications such as in drug response prediction (24, 29). VAEs have been able to generate biologically-relevant data embed-dings (25, 30) and offer flexibility in generating synthetic data by directly manipulating them (31). In this work, we propose a methodology based on the VAE to generate synthetic microarray instances of MB patients robustly and in an explainable way (summarized in Figure 1).

**Figure 1.**
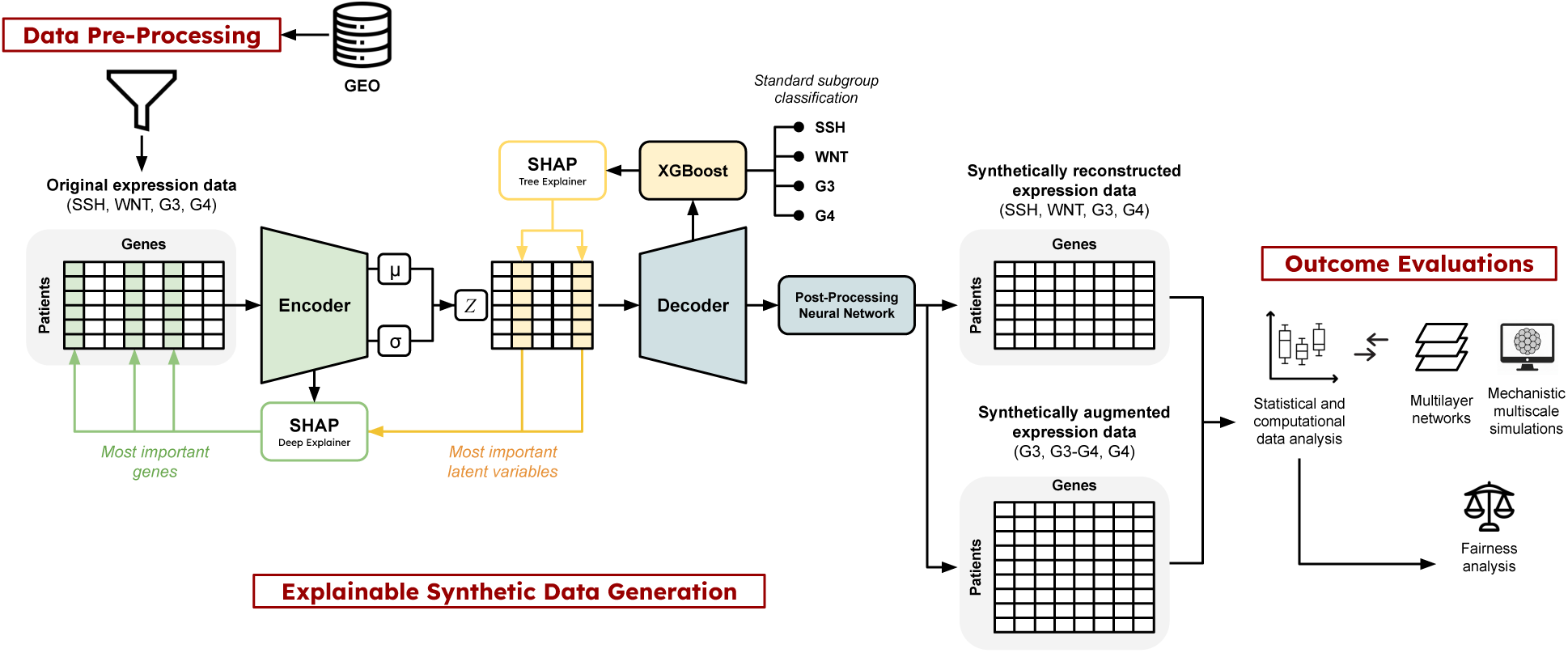
Overview of the workflow of the study. The data for this study was obtained from a previous publication (10), that made it accessible on GEO. The data was preprocessed to remove low-variance and outlier genes, to then train the VAE. The VAE consists of two fundamental parts. First, the encoder reduces the input dimension to an arbitrary number. For each of the reduced dimensions, two parameters are obtained, representing the mean and standard deviation of a Normal distribution, making up the latent space. Second, a decoder recovers the original space of the data, with an associated reconstruction error. In order to minimize this error, the output of the decoder is used to train a post processing neural network. Using the generative ability of the decoder, it is possible to create synthetic data that resembles the original, and use it to study the subgroups of MB with high specificity. Besides, in order to explain the relationship between MB subgroups and genes, a classifier was used on the latent space to distinguish the four MB subgroups (SHH, WNT, Group 3, and Group 4). This classification may be explained by obtaining the genes’ SHAP values, which requires two steps. SHAP’s TreeExplainer obtains the relationship between the classified subgroups and the latent space. The most important latent variables (those that explain most of the classification) are then passed to SHAP’s DeepExplainer to obtain the genes that explain the subgroup classification.

Here, we aim to test the hypothesis that an intermediate group with consistent characteristics may exist between G3 and G4, building on our previous findings using independent multi-omics data and complementary methodology. We develop an unsupervised methodology to identify a set of patients with uncertain clustering, i.e., patients that are not clearly in either G3 or G4. We then use the generative capabilities of the VAE to generate new, synthetic patients within this identified cluster and apply a pipeline based on SHapley Additive exPlanations (SHAP) (32) to the VAE to better understand the link between the learned latent space’s dimensions and the input gene expression levels. Moreover, differential expression analysis reveals significant differences in gene expression between G3, G4, and the identified subgroup G3-G4.

In this regard, model explainability and confidence of our findings about the identified subgroup are critical aspects of our work. While our SHAP-based pipeline addresses model explainability, the confidence in the validity of our results is supported by comparing our results with previous studies. In particular, a G3-G4 cluster was identified in our previous study using a different dataset and methodology, based on multilayer networks, revealing a remarkable overlap in molecular characteristics among the patients identified independently (14).

A key contribution of this work is showing that overlooking the intermediate G3–G4 subgroup may result in treatment disparities, as patients from this cluster could be misclassified and consequently receive overly aggressive therapy intended for G3 or insufficient treatment designed for G4. Such use of Machine Learning Fairness approaches on synthetic data opens new opportunities for research, especially in rare disease contexts. Beyond refining subgroup boundaries with VAEs and synthetic cohorts, it is also important to test whether the inferred G3, G3-G4 and G4 classes also differ in their signaling pathways activities. To this end, we used the same real and synthetic samples to personalize a curated Boolean model of Group 3/4 medulloblastoma and derived pathway-level activation profiles from MaBoSS stationary distributions, thereby asking whether the intermediate G3-G4 subgroup is also distinguished at the level of oncogenic pathway usage rather than only by gene expression.

Overall, our results present the potential of SDG pipelines, particularly explainable VAE-based deep learning models, to better understand the underlying biology of the more complex and heterogeneous MB molecular subgroups. Thanks to synthetic data generation, this approach is well suited for research with limited or highly specific data, such as rare diseases or underrepresented patient groups, and enables the evaluation and mitigation of potential inequities.

## 2. Results

### 2.1. Identifying an Intermediate Subgroup Between G3 and G4

We analyzed RNA microarray data in GEO from 763 primary medulloblastoma samples, obtained from the largest transcriptomics study available to date on medulloblastoma (MB) (10). After preprocessing (Methods 4.1), the dataset comprised 14,349 genes across 763 patients.

The dataset consists of samples labeled to the four classical subgroups (G3, G4, SSH and WNT), with a noticeable group imbalance (Table 1). To get a first glimpse of the separability of the groups, we visualized the data using a UMAP (33) representation (Figure 2A). The classification of the four subgroups using a weighted XGBoost (34) achieved high performance (Figure 2D). Nevertheless, as expected from the literature (see Introduction), a less clear separability is observable between G3 and G4 from the partial overlap that can be seen in the UMAP visualizations, and from clustering results (Supplementary Figure S8A). We hence focused on G3 and G4 patients.

**Figure 2.**
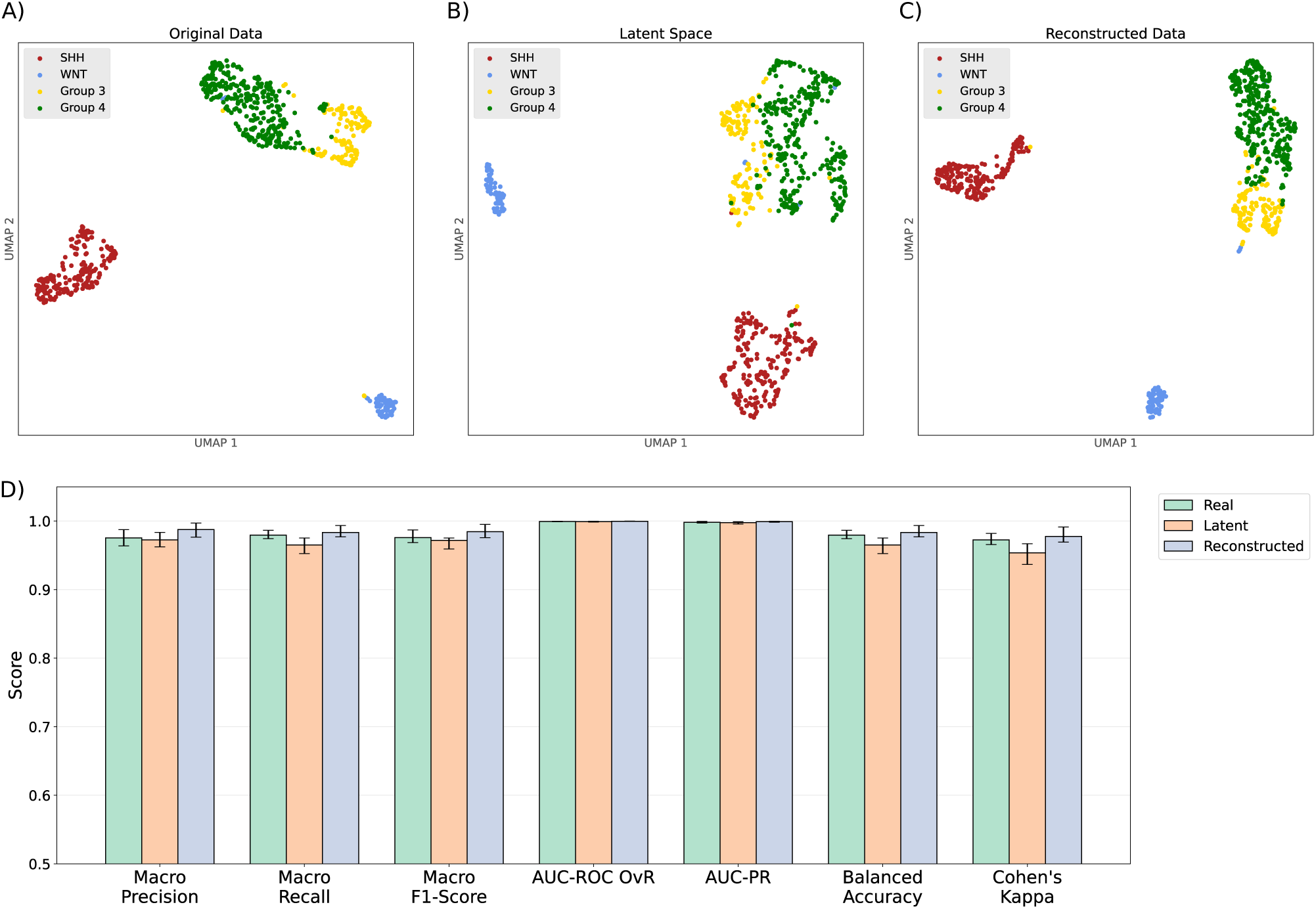
Exploration of the four medulloblastoma subgroups across the original and learned data spaces. A-C) UMAP of the subgroups. The four subgroups (SHH, WNT, Group 3, and Group 4) are highlighted in different colors. The UMAP reflects the clear separability of the SHH and WNT subgroups from each other and a third cluster, which represents Groups 3 and 4. In this latter cluster, some patients are not clearly separated but overlapping. As can be appreciated, the learned representations resemble the original data UMAP. **D) Classification performances for the subgroups in the real, latent space, and reconstructed spaces.** Figures showing the classification results for the four medulloblastoma subgroups in the different data spaces considered in this study. Evaluation metrics include macro precision, macro recall, macro F1-score, area under the ROC curve (AUC-ROC) using the One-vs-Rest (OvR) approach (AUC-ROC OvR), area under the precision-recall curve (AUC-PR), balanced accuracy, and Cohen’s Kappa. The values shown represent the median scores from 10 runs, each with a different random seed. Each bar includes whiskers extending from the 25% to the 75% score quantile. The median for each classification and metric are stated in Table 2

**Table 2.**
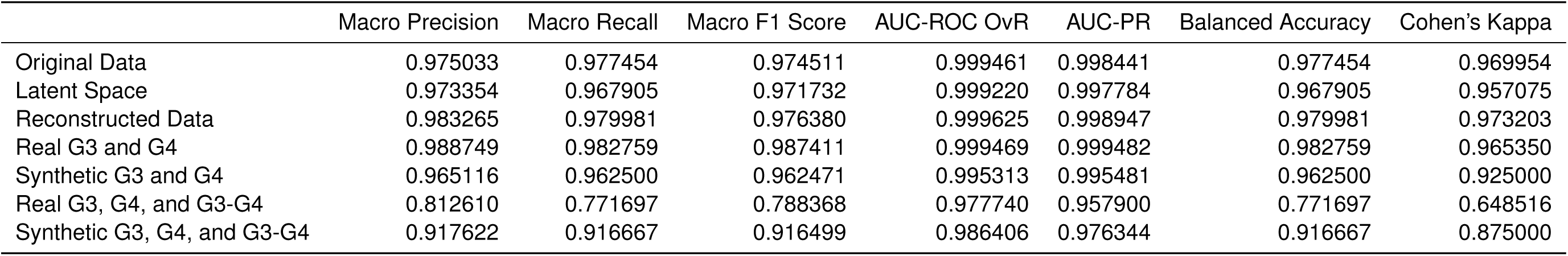
Classification results. This table shows the values for the median results of all the classifications performed in the present work, each repeated 10 times with different, random seeds. Evaluation metrics include macro precision, macro recall, macro F1-score, area under the ROC curve (AUC-ROC) using the One-vs-Rest (OvR) approach (AUC-ROC OvR), area under the precision-recall curve (AUC-PR), balanced accuracy, and Cohen’s Kappa.

|  | Macro Precision | Macro Recall | Macro F1 Score | AUC-ROC OvR | AUC-PR | Balanced Accuracy | Cohen's Kappa |
| --- | --- | --- | --- | --- | --- | --- | --- |
| Original Data | 0.975033 | 0.977454 | 0.974511 | 0.999461 | 0.998441 | 0.977454 | 0.969954 |
| Latent Space | 0.973354 | 0.967905 | 0.971732 | 0.999220 | 0.997784 | 0.967905 | 0.957075 |
| Reconstructed Data | 0.983265 | 0.979981 | 0.976380 | 0.999625 | 0.998947 | 0.979981 | 0.973203 |
| Real G3 and G4 | 0.988749 | 0.982759 | 0.987411 | 0.999469 | 0.999482 | 0.982759 | 0.965350 |
| Synthetic G3 and G4 | 0.965116 | 0.962500 | 0.962471 | 0.995313 | 0.995481 | 0.962500 | 0.925000 |
| Real G3, G4, and G3-G4 | 0.812610 | 0.771697 | 0.788368 | 0.977740 | 0.957900 | 0.771697 | 0.648516 |
| Synthetic G3, G4, and G3-G4 | 0.917622 | 0.916667 | 0.916499 | 0.986406 | 0.976344 | 0.916667 | 0.875000 |

**Table 3.** p-values, effect sizes and 95% confidence interval (CI) of the effect size for the comparisons between the three groups among the commonly mutated genes mentioned in. (**45**). The table shows the statistics corresponding to the DEA among the three groups (G3, G4, G3-G4) for the genes that are commonly mutated in medulloblastoma according to (45).

|  | Group 3 vs Group 4 |  |  | Group 3 vs G3-G4 |  |  | Group 4 vs G3-G4 |  |  |
| --- | --- | --- | --- | --- | --- | --- | --- | --- | --- |
|  | p-value | Effect Size | 95% CI Effect Size | p-value | Effect Size | 95% CI Effect Size | p-value | Effect Size | 95% CI Effect Size |
| KMT2C | 4.75e-04 | -0.397 | [-0.595,-0.198] | 2.25e-04 | 0.398 | [0.199,0.596] | 3.05e-14 | 0.851 | [0.645,1.056] |
| MYC | 4.07e-44 | 1.715 | [1.485,1.945] | 2.60e-06 | 0.638 | [0.436,0.839] | 2.96e-19 | -1.106 | [-1.317,-0.895] |
| SNCAIP | 4.89e-46 | -1.780 | [-2.012,-1.547] | 5.07e-18 | -1.106 | [-1.317,-0.895] | 8.85e-08 | 0.644 | [0.442,0.846] |
| SYNCRIP | 1.62e-04 | 0.407 | [0.209,0.606] | 1.24e-22 | 1.091 | [0.88,1.301] | 1.36e-08 | 0.641 | [0.44,0.843] |
| TP53 | 2.01e-17 | 0.920 | [0.714,1.127] | 6.03e-06 | 0.527 | [0.327,0.727] | 3.29e-04 | -0.412 | [-0.611,-0.214] |

**Table 4.** p-values, effect sizes and 95% confidence interval (CI) of the effect size for the comparisons between the three groups among the commonly mutated genes mentioned in. (**14**). The table shows the statistics corresponding to the DEA among the three groups (G3, G4, G3-G4) for the genes that were also found to be differentially expressed by (14).

|  | Group 3 vs Group 4 |  |  | Group 3 vs G3-G4 |  |  | Group 4 vs G3-G4 |  |  |
| --- | --- | --- | --- | --- | --- | --- | --- | --- | --- |
|  | p-value | Effect Size | 95% CI Effect Size | p-value | Effect Size | 95% CI Effect Size | p-value | Effect Size | 95% CI Effect Size |
| CDKN2C | 6.76e-07 | 0.549 | [0.349,0.749] | 7.99e-22 | 1.031 | [0.822,1.24] | 1.72e-05 | 0.471 | [0.271,0.67] |
| IMPDH1 | 5.29e-25 | 1.145 | [0.933,1.357] | 2.68e-11 | 0.786 | [0.582,0.99] | 9.19e-04 | -0.426 | [-0.625,-0.227] |
| SLC35A4 | 2.30e-28 | 1.199 | [0.985,1.412] | 1.94e-10 | 0.762 | [0.559,0.966] | 1.20e-05 | -0.422 | [-0.621,-0.223] |
| ST8SIA2 | 2.69e-02 | -0.280 | [-0.478,-0.083] | 8.86e-05 | 0.298 | [0.1,0.495] | 3.35e-11 | 0.690 | [0.487,0.892] |
| ST8SIA3 | 3.10e-12 | -0.738 | [-0.941,-0.535] | 1.81e-02 | 0.090 | [-0.106,0.287] | 1.66e-22 | 1.037 | [0.828,1.247] |

To identify patients with shared characteristics with G3 and G4, we attempted to reproduce the methodology proposed in (10) based in consensus clustering, for which no code is available. Due to lack of information, we performed the analysis both on the unmodified data downloaded from GEO and on our preprocessed data. In both cases, the number of patients that we observed (Supplementary Figure S2A,B, and Methods 4.1) were inconsistent compared to those reported in (10). Given the limitations of this procedure proposed in (10), we developed a new methodology to more robustly obtain patients belonging to the G3-G4 subgroup.

Our approach consists in a k-Nearest Neighbors Graph (k-NNG) followed by an agglomerative clustering to discover the optimal number of neighbors in the data from G3 and G4 (see Methods 4.2). Supplementary Figure S8A shows the silhouette score for different numbers of neighbors *k*, with a maximum at *k* = 2 and 2 clusters (one cluster with 341 patients and the other with 129). Noticeably, when choosing 3 clusters very similar numbers are achieved on average, pointing out that a division into three clusters may be possible (with *k* = 2, the three clusters had 247, 129, and 94 patients). We fixed the number of neighbors to 2 and implemented an analysis to determine those patients not clearly assigned to either G3 or G4. To do so, we repeated the clustering 1,000 times, each time randomly sampling 100 patients from each subgroup. We counted how many times a patient was assigned to a different cluster when compared to the clinical metadata. The result is a distribution of the number of times a patient was assigned to a different cluster (Supplementary Figure S8B,C). The cutoff to determine which patients should be assigned to the G3-G4 subgroup was set to the expected value that patients, chosen at random, would change subgroups (formulas included in Methods 4.2). These top cluster-changing patients, above the respective cutoffs, are considered of ‘uncertain label’ and shown in Supplementary Figure S2C. Supplementary Table S2 shows the number of patients in the G3-G4 subgroup and which subgroups they originally belonged to. Although most detected G3-G4 samples had originally been labeled as G4 in (10), a two-tailed a Mann-Whitney U on the age distribution between them was significantly different (U = 3664.5, p = 0.009, Cohen’s d = 0.496), as seen in Supplementary Figure S3. However, no difference was found between G3 and G3-G4 (U = 1003.5, p = 0.249, Cohen’s d = 0.039).

In (10), 14 G3-G4 patients were found, but these were not given labels. In our case, the number of patients in the G3-G4 subgroup that we identified is also small, namely 19 patients (in contrast, Supplementary Figure S4 shows the results of label-randomization experiments, where a median of 262 patients are found to be intermediate), compared to the number of patients in other subgroups (Supplementary Table S2), so any numerical analysis may be of limited statistical significance. Therefore, we sought to augment the data within the G3-G4 subgroup to balance it with the other subgroups and better study its molecular characteristics.

### 2.2. A Pipeline to Generate Reliable Synthetic Data of Rare Disease with the VAE

Having identified the samples belonging to the G3-G4 subgroup, we then resorted to a generative modeling approach to create synthetic data that could help us better describe the characteristics of the G3-G4 subgroup. In order to create synthetic data, we leveraged the ability of the Variational Autoencoder (VAE) to learn a lower-dimensional representation of the data, or latent space, where new instances can be sampled from and decoded back to the original space.

We trained a VAE on the preprocessed RNA microarray dataset. The architecture of the VAE (shown in Supplementary Figure S1) consists of an encoder, which learns to represent the data into a latent space, and a decoder, which learns to reconstruct the data from the latent space. The encoder and the decoder are modeled as neural networks, each with two hidden layers. We explored various configurations of hidden layers and latent space dimensions, identifying the optimal configurations using the Mean Squared Error (MSE) between the real and the generated data (Supplementary Figure S5A; Supplementary Table S1; see Methods 4.3). We chose the model with the lowest MSE, corresponding to a hidden layer dimension of 1,024 neurons and a latent space of 64 dimensions. Additionally, we trained an independent neural network to post-process the VAE outputs, consistent with multi-stage approaches where a dedicated model is used to refine and improve the fidelity of VAE-generated results (e.g. in (35)) (see Methods 4.4). The suitability of this model was further confirmed by the inspection of the reconstructed data using the Wasserstein distance between the real and the generated data (Supplementary Figure S7). Overall, thanks to the post processing neural network, the reconstruction error was reduced from 26.8 to 0.04 on average for the last 20 test epochs (Supplementary Figure S6). The objective of the added network is to act as a refinement step, aligning synthetically-generated samples with the empirical data manifold to ensure compatibility in downstream tasks.

We further validated the suitability of our VAE model by evaluating the classification of medulloblastoma subgroups in the original, latent and reconstructed spaces (Figure 2D). Medians for each performance metric is detailed in Table 2. The UMAP visualization of the original, latent and reconstructed spaces are shown in Figure 2A-C reflecting the separability between the subgroups and possible overlap between G3 and G4. Furthermore, a 3D PCA visualization of the latent space in Figure 3 places the putative G3-G4 intermediate subgroup at the boundary between G3 and G4.

**Figure 3.**
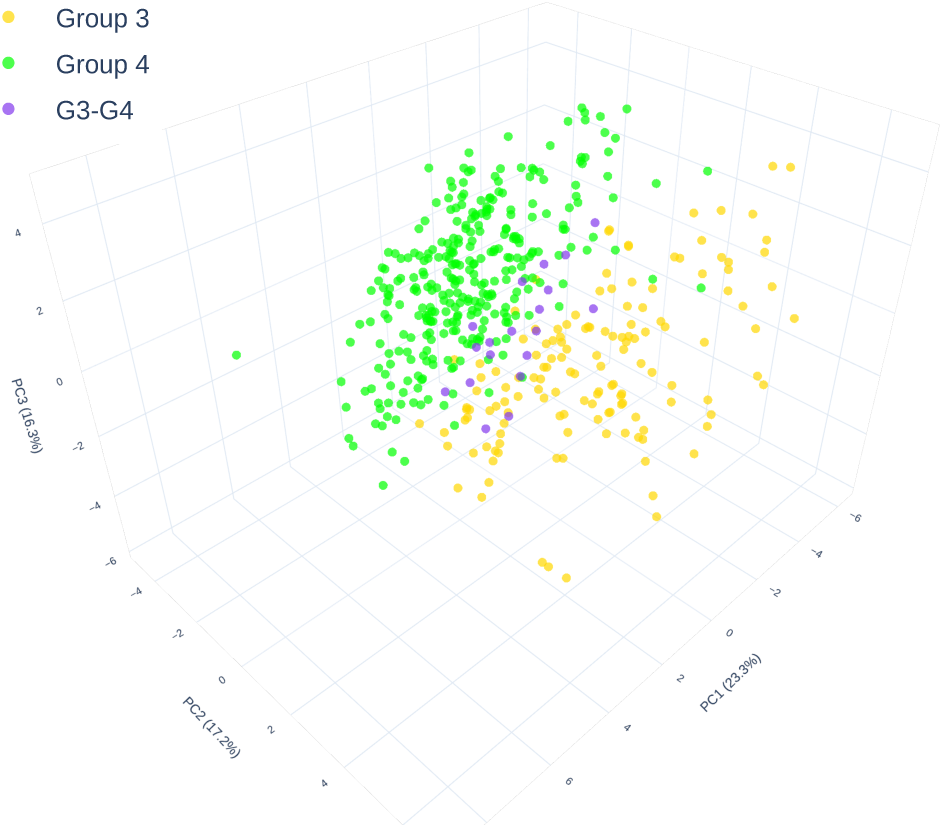
PCA of the VAE latent space, with the intermediate G3-G4 subgroup identified. The 3-dimensional PCA shows the spread of the dataset in the VAE latent space, showing all the real patients of the labeled Group 3 (G3) and Group 4 (G4) patient samples. At the intersection of both groups are situated the identified samples belonging to the putative intermediate subgroup, G3-G4. Each of the PCA’s axis includes the percentage of variability explained by the corresponding principal component (23.3%, 17.2%, and 16.3%). PC: Principal Component.

To further test the reliability of the synthetic data generation, we compared two strategies for generating synthetic instances within each subgroup. First, we added random noise to a sampled set of the real data. Second, we sampled patients in the latent space, added noise, and decoded them back to the original space. Both of these methodologies are agnostic to the patient labels. Namely, with our synthetic data generation pipeline, synthetic data can be produced in any region of the latent space, which captured the statistical characteristics of the real data points of interest. Figure 4A-D shows the UMAP projection in the real and reconstructed spaces for varying noise levels. As can be appreciated, slight noise levels in the real and latent space yield similar patients to those in the original data However, as the noise level increases, synthetic patients generated in the real space differ from the original patients (Supplementary Table S4). We chose a noise level of 0.2 because it can generate synthetic patients that are different from the original ones, while still keeping the subgroup separability. UMAPs with larger noise levels are reported in Supplementary Figure S9. That is not the case for the synthetic patients generated in the latent space and then reconstructed, which remain similar to the original patients and keep subgroup separability. This indicates that the VAE can reliably generate meaningful synthetic data that highly resembles the real data when adding noise to the latent representations. The effectiveness of adding noise in the latent space can be attributed to the smoothness and lower dimensionality of the latent representation, which allows for controlled and meaningful perturbations aligned with the learned data distribution, a property that has been recently leveraged by more sophisticated approaches in different domains (36).

**Figure 4.**
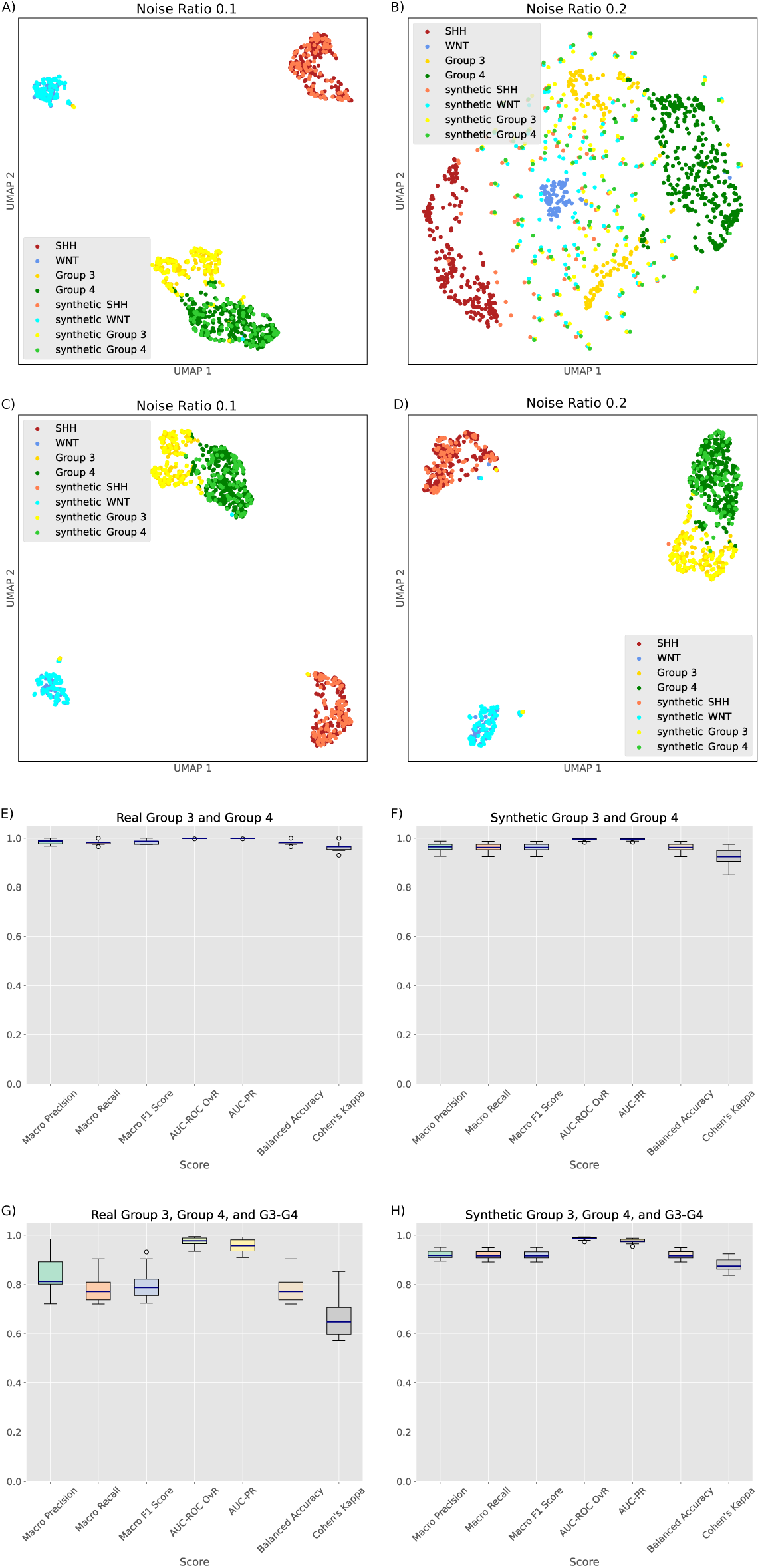
Exploration of the synthetic data. A-D) UMAPs comparing a control method for data generation with our VAE-assisted method. A,B) show the UMAPs of the real data augmented with Gaussian noise. C,D) show the UMAPs of the synthetic data, which was augmented with noise in the latent space, then reconstructed with our pipeline. The sub-panels are organized by columns, representing increasing levels of added noise. These results demonstrate that adding noise directly to real patient data produces nonsensical outputs. However, when using the patients’ latent representation and the VAE’s decoder, increasing levels of noise yield increasingly different, yet similarly distributed synthetic patients. **E-H) Classification results**. The cases considered are: E) real Groups 3 and 4, F) synthetic Groups 3, 4 G) real Groups 3, 4, and G3-G4, H) synthetic Groups 3, 4, and G3-G4. E) and F) reflect the appropriateness of our synthetic-data-generation pipeline, which produces patients that resemble the original separability. G) and H) show how our synthetic data generation, used to balance out the three groups of interest, results in a better separability when the putative G3-G4 subgroup is considered.

Based on this result, we augmented the number of patients in this subgroup to 200 (Methods 4.4) by adding a noise ratio of 0.2 to the real patients in the latent space. The resulting latent representations were then decoded, producing the corresponding synthetic patients in the real data space. Analogously, we used our pipeline to balance out the other two subgroups of interest: Groups 3 and 4. The UMAP of the real data overlaid with 200 synthetic patients for each of the three subgroups of interest, is shown in Supplementary Figure S10. As a result of our reconstruction pipeline, most synthetic patients appear alongside their real counterparts. The UMAP visualization appears to separate Groups 3 and 4, although the UMAP representation may not be a trustworthy indicator of global data structure (37). To further test this separability, we ran another set of classifications, whose results are shown in Figure 4E-H and Table 2. Figure 4E shows the classification result of real patients in Groups 3 and 4, while Figure 4F shows the result of classifying synthetic patients of Groups 3 and 4; confirming the separability of both real and synthetic cases is similar. When adding the G3-G4 subgroup to the real case (Figure 4G), performances drop significantly. However, when balancing the data with the generated synthetic instances, very high performances are recovered (Figure 4H). The consistently high classification performance (Table 2) in the synthetic balanced scenario when G3-G4 is introduced (only 1.9% difference in AUC-PR and reduced variance across folds) supports the definition of three distinct groups, while mitigating concerns about possible bias in the two-group classification due to the underrepresentation of a potential third group. This reflects a common issue in classification involving potential mislabeling of rare occurrences, which is often overlooked in biomedical data analysis but can lead to serious consequences (38, 39, 40). For example, in the context of MB, G3 is typically more aggressive than G4 (41), and misclassifying G3-G4 patients could lead to inappropriate treatments, as previously highlighted (42, 43).

### 2.3. The Augmented G3-G4 Subgroup Reveals Unique, Differentially Expressed Genes

To achieve a better understanding of the molecular components defining the G3-G4 subgroup, we identified genes with differential transcriptomic expression across G3, G4 and G3-G4, which had been amplified and balanced out using the VAE. We ran the analysis using limma (44), which returned 11,869 differentially expressed genes between the three subgroups (Supplementary Dataset 1). Given the large number of genes identified, we looked for a more stringent method. To this end, a non-parametric Kruskal-Wallis test (p<0.01) was performed on the three compared subgroups, followed by Dunn’s test with Bonferroni adjustment (p<0.01), to find out which genes showed significant differences in the median between the subgroups of interest (see Methods 4.5). Supplementary Dataset 2 contains all significantly differentially expressed genes along with the corresponding p-values when comparing the three groups with Dunn’s test, the effect size and the corresponding 95% confidence intervals for the effect size. 2,506 genes were found to be significantly differentially expressed between the synthetic patients in the three subgroups. When we calculated the effect size, 757 genes showed a substantial effect size (Cohen’s d > 0.5) across the three subgroups, and 120 genes showed a very pronounced effect size (Cohen’s d > 0.8).

We then selected genes that previous studies (45) found to be commonly mutated in each subgroup (Table S3). As a result, we encountered several relevant genes that showed significant differences between the subgroups: KMT2C, MYC, SNCAIP, SYNCRIP, and TP53 (Figure 5A-E and corresponding p-values and effect sizes on Supplementary Table 3). Mutations in these genes are known to have significant impacts in pediatric brain cancer development and progression. MYC family genes c-MYC and MYCN amplifications are associated with G3 poor prognosis, and observed in SHH and G4, respectively (46). TP53 mutations are an important risk factor in SHH (47). KMT2C mutations in pediatric brain tumors can alter chromatin structure and serve as a potential biomarker for immunotherapy response (48). SNCAIP alterations serve as a novel diagnostic biomarker for G4, where they can disrupt the local chromatin environment (49). Although not specifically mentioned in pediatric brain tumors, SYNCRIP mutations in other cancers promote resistance to targeted therapies and contribute to tumor heterogeneity through increased mutagenesis (46).

**Figure 5.**
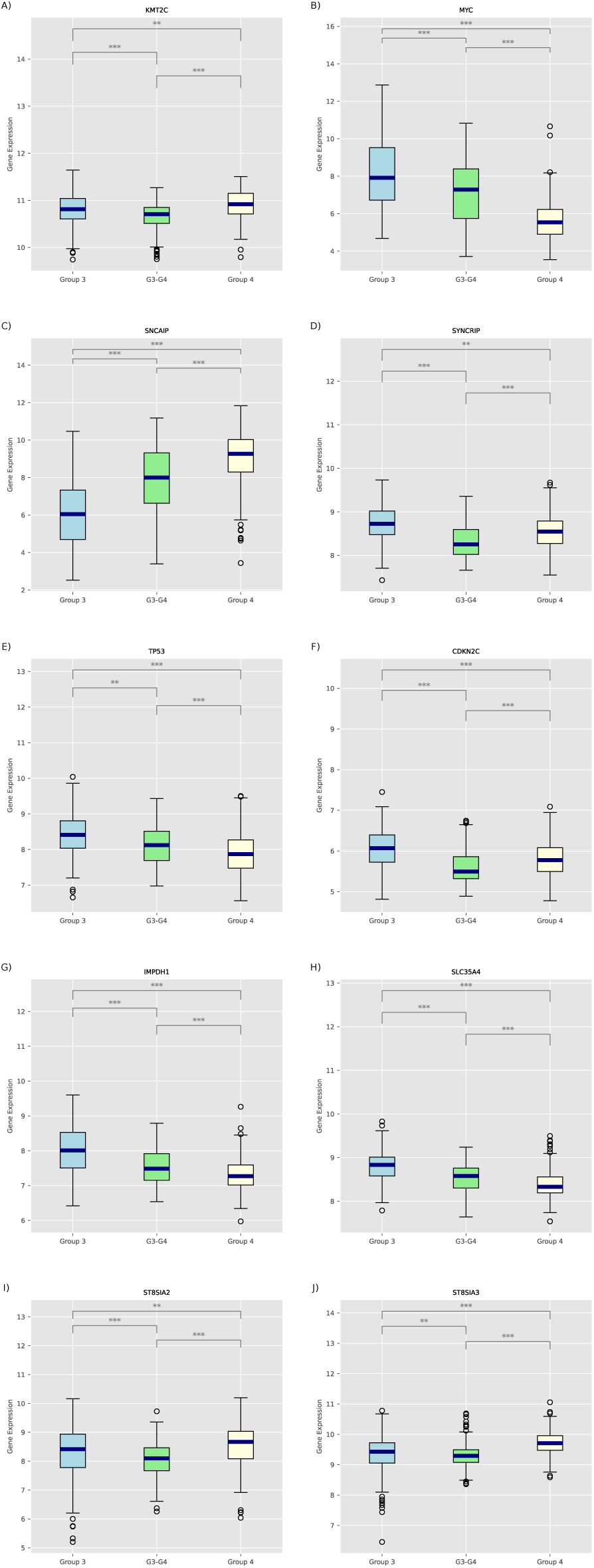
Identifying relevant genes within MB subgroups. A-E) Differentially expressed genes in the G3-G4 subgroup compared to Groups 3 and 4 that are known to be commonly mutated in medulloblastoma. KMT2C, MYC, SNCAIP, SYNCRIP, and TP53. These genes are commonly mutated in medulloblastoma, according to (45). **F-J) Genes that our previous publication found to be important to determine the G3-G4 subgroup.** CDKN2C, IMPDH1, SLC35A4, ST8SIA2, ST8SIA3. Differentially expressed genes in accordance with our previous study (14), that also detected patients as part of a putative G3-G4 subgroup. All panels show the differentially expressed genes between synthetic data of Groups 3, 4, and the G3-G4 subgroup. These genes were found to be differentially expressed between the subgroups through a Kruskal-Wallis and a Dunn’s test for the median. The boxplot shows the distribution of the gene expression values for each gene and subgroup. The central line represents the median, the box the interquartile range, the whiskers 1.5 times the interquartile range, and the white dots the outliers. Notice the y-axis scale is different for each plot. Horizontal lines with asterisks indicate statistically significant differences between groups. Asterisks denote the level of significance: **p < 0.01, ***p < 0.001.

Given the diffuse definition of the existing continuum between G3 and G4 in the literature (50), we aimed to compare the molecular features of our proposed G3-G4 subgroup with that of a previous publication that identified three patients as belonging to the potential G3-G4 subgroup using multi-omics data and a different methodology (14). Among the differentially expressed genes within the G3-G4 subgroup, we observed: CDKN2C, IMPDH1, SLC35A4, ST8SIA2, and ST8SIA3 (Figure 5F-J and corresponding p-values and effect sizes on Supp. Table 5). CDKN2C, a member of the INK4 family of cyclin-dependent kinase inhibitors, is a gene known for its involvement in MB33. IMPDH1 has been previously associated with a worse prognosis in glioma (51). ST8SIA2 was among the highest-expressed upon mice treatment with digoxin, a chemical recently described as a potential treatment for both Groups 3 and 4 (52). ST8SIA3 is a paralog of ST8IA2, also described in (52). Although to our knowledge, SLC35A4 has not been previously related to MB, members of the SLC35 family, including SLC35A2, SLC35A3, and SLC35A1, have been associated with neurological conditions such as early-onset epileptic encephalopathy (53). A comparison of the expression levels of each gene for the 3 subgroups of interest is shown in Figure 5F-J.

### 2.4. Subgroup Classification in Latent Space is Explained by a Distinct Set of Genes

The genes we identified provide insights into the molecular differences between G3-G4, G3, and G4, but equally important is explaining how our developed VAE models MB. Deep Learning models, such as the VAE, are often considered black-box models. Therefore, there is a challenge when providing a clear biological understanding of the relationship between the input genes and the inferred latent space. This lack of interpretability is deemed a significant drawback for the application of such models in clinical settings (54). Hence, we aimed to gain an understanding of the key genes that explain the separation between the four original medulloblastoma subgroups—i.e., we wanted to understand how the VAE can model the MB disease globally, as opposed to the specific case of the G3-G4 subgroup we had determined earlier.

To gain such knowledge about MB, we calculated SHapley Additive exPlanations (SHAP) values (32) in a two-step approach. We first classified the medulloblastoma subgroups in the latent space using an XGBoost model, optimized for hyperparameters, and then interpreted the results with SHAP’s TreeExplainer to uncover the relationship between subgroups and latent variables (see Methods 4.6). SHAP values allow us to identify the most important latent variables as those that showed an absolute SHAP value larger than average (Supplementary Figure S11A). We then used SHAP’s DeepExplainer to analyze the importance of genes in generating those most important latent variables, effectively disentangling the relationship between the latent variables and the gene expression variables (see Methods 4.6). Supplementary Figure S11B shows the distribution of the genes’ SHAP values. We chose a the top 5% of the distribution, or 717 genes, as a non-parametric criteria for ranking the most important genes given the non-normality of the SHAP values distribution. A 1% threshold was tested but resulted in some inconclusive analyses from gprofiler (Supplementary Figure S13). A full list of these genes is available in the Supplementary Dataset 3. Among them, we highlight the presence of MYC, SNCAIP, and ST8SIA2, which had already appeared in the previous section (see Results 2.3). A functional enrichment analysis with *gprofiler* (55) (Methods 4.7) on the 717 genes reveals the most common biological functions are related to the neural system and development (Supplementary Dataset 3 and Supplementary Figure S12), which had been linked to G3 and G4 (12).

### 2.5. Mechanistic Boolean Models Allows For Subgroup Classification

Having established, using VAEs and synthetic data generation, that a transcriptionally intermediate G3-G4 subgroup improves both subgroup separability and classifier fairness, we next asked whether these three classes also differ in their underlying signaling architecture. We therefore used 763 patients (10) and 600 synthetic samples from this work to personalize a manually curated Boolean model of Group 3/4 medulloblastoma signaling, encompassing Proliferation, Apoptosis, PI3K-AKT-mTOR, RAS-MAPK, Hippo-YAP, RHO-GTPases, and Motility modules (see Methods 4.9), and used MaBoSS a Boolean simulator designed to estimate per-sample stationary activation probabilities for all nodes.

Using the activation probability of each node from MaBoSS, we grouped them in the aforementioned pathways, averaged them and compared these mean activation probability scores between groups (Figure 6). This clustered heatmap reveals systematic subgroup-specific differences in pathway usage that closely mirror the transcriptomic stratification. Notably, the real and synthetic groups clustered together for each group, strengthening the previous results of this study, and different pathways had significantly different activities across all groups (Kruskal-Wallis test, p < 0.05). The pathway activity profiles derived from the simulations across G3, G4, and intermediate G3-G4 medulloblastoma states delineate a coherent signaling continuum that is consistent with the literature (Figure 6, Supplementary material). In our simulations, G3 and the corresponding synthetic G3 is characterized by the highest proliferation and cell-cycle scores, driven by MYC and CDC25B, together with comparatively lower PI3K-AKT-mTOR and cytoskeleton/motility activities (Mann-Whitney-Wilcoxon pairwise tests comparisons and Cohen’s d effect size tests available in Supplementary material). In contrast, Group 4 and synthetic G4 displayed reduced proliferation but markedly increased PI3K-AKT-mTOR activation and the strongest engagement of Rho-GTPase-dependent cytoskeletal remodeling and motility/invasion modules, including SRC and CDC42/RAC1/RHOA-PAK-ROCK signaling. Intermediate G3-G4 and synthetic G3-G4 occupy transitional positions for these pathways, with proliferation and survival signaling lying between G3- and G4-like extremes and increased RTK activity. Thus, the Boolean model confirms that the VAE-defined G3-G4 subgroup is not a statistical artifact of the latent space, but sits at an intermediate position at the biological functional level along a continuum from MYC-high, highly proliferative Group 3 signaling to RTK/PI3K-AKT-mTOR- and motility-dominated Group 4 signaling.

**Figure 6.**
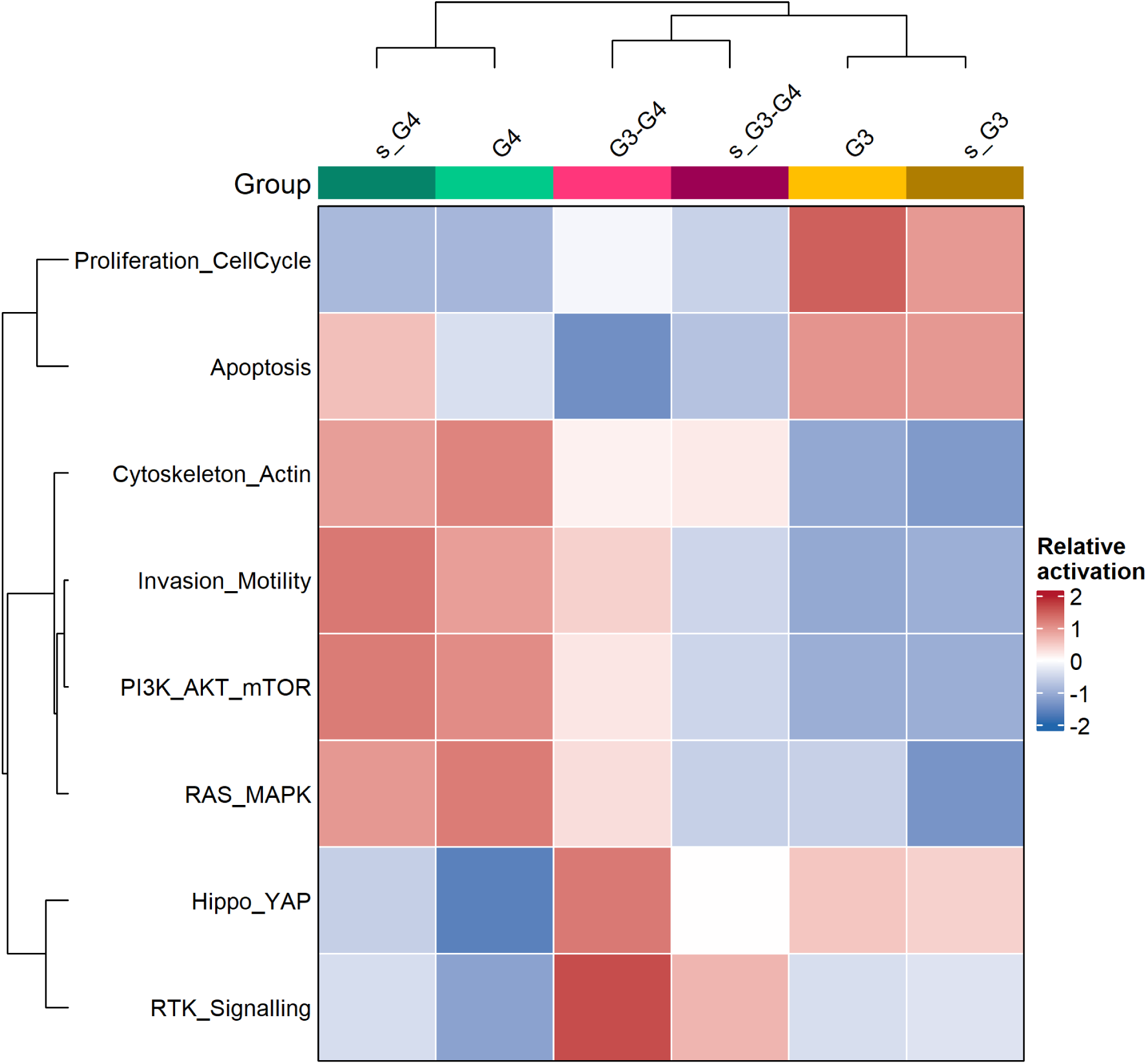
Heatmap of pathway-level Boolean model activation profiles across medulloblastoma groups. Each column represents one patient group (Group 3, Group 3-Group 4, Group 4 and their synthetic counterparts), and each row corresponds to a pathway score obtained by averaging the activation probabilities of all Boolean model variables assigned to that pathway. Mean pathway scores were first computed per group, then standardized per pathway (row-wise z-scores), and visualized using the Euclidean distance and complete-linkage clustering on rows and columns. Colors indicate relative activation within each pathway (blue: below-average, white: average, red: above-average; scale in z-score units), with the top annotation denoting group identity.

Additionally, these simulation-based patterns integrate well with the literature, where G3 tumors are more strongly defined by MYC-associated cell cycle and translational control (46, 1), and aberrant SRC signaling and downstream MAPK, PI3K, and mTOR activation constitute a hallmark of G4 medulloblastoma (15). The simulation results recapitulate this division at the level of pathway activities: G3 correspond to a high MYC activity, highly proliferative but relatively RTK/PI3K-mTOR-low configurations, while G4 correspond to more differentiated, survivaland motility-oriented configurations with enhanced PI3K-AKT-mTOR and cytoskeletal signaling, and G3-G4 lay in between both, with a notable RTK activity.

Furthermore, the existence and positioning of the intermediate G3-G4 states in our simulations are compatible with a recent work that resolve a multi-axial Group 3/4 landscape composed of photoreceptor-like (PR), MYC-enriched, precursor-like and unipolar brush cell-like (UBC) gene regulation axes along the cerebellar UBC lineage (56). Group 3 tumors are enriched for PR/MYC states, whereas Group 4 tumors are enriched for Precursor/UBC states; mixed subtypes (including those with intermediate Group 3/4 scores) contain coexisting PR and UBC trajectories emerging from a shared precursor pool. Within this framework, our G3 and synthetic G3 correspond to earlier MYC-axisdominated progenitor states; G4 and synthetic G4 correspond to more UBC-like, RTK/PI3K-mTOR- and motilitydriven states; and G3-G4/synthetic G3-G4 represent coarse-grained mixtures near the bifurcation between these trajectories.

### 2.6. Only the Balanced Synthetic Data Multi-Class Classifier Achieves Fairness Across Groups

Despite consistently larger variances in predictive metrics, the binary classifier trained on balanced synthetic data (henceforth, 2SD) shows slightly better overall performance than the multi-class classifier trained on balanced synthetic data (henceforth, 3SD) (see Figure 4). Beyond overall performance, however, a key question is whether classification accuracy is achieved equally across groups, i.e., whether the assigned labels reflect a fair label assignment rather than systematically favoring certain groups. As a fairness criterion, we use Equalized Odds (57), requiring equal True Positive Rates (TPR), or Equal Opportunity, and equal False Positive Rates (FPR), or Predictive Equality, across groups. This ML fairness notion is particularly important in sensitive fields like healthcare, criminal justice, and hiring (58). Given high accuracy does not guarantee fair outcomes across groups, unfair models should be discarded, as supported by many recent publications and best practices (59, 60, 61). To assess this, we evaluated all classifiers, trained on real data (2RD, 3RD) and synthetic data (2SD, 3SD), based on the Equalized Odds criterion.

As our fairness comparison encompasses both binary and multi-class scenarios, we used weighted macro TPR and weighted macro FPR, which accounts for class imbalances by adjusting for class support (62) (see Methods). This approach provides a consistent way to evaluate group-level fairness across different model types.

Also, to prevent any data leakage, all classifiers were trained and evaluated using Leave-One-Out Cross-Validation (LOOCV), which ensures unbiased performance estimates (63), with hyperparameters optimized within each fold (see Methods).

For this fairness assessment, we defined a baseline tri-part division of patients: the “Intermediate group” (19 patients), previously identified in an unsupervised way (see Results, “Identifying an Intermediate Subgroup Between G3 and G4”), and the remaining “More aggressive group” (142 patients) and the “Less aggressive group” (309 patients), associated with metadata provided by Cavalli’s study (10). A fair classifier, whether binary (using labels G3 and G4 as ground truth) or multi-class (using labels G3, G4, and G3–G4 as ground truth), should achieve comparable detection and error rates across the three groups.

Comparing each classifier’s Equal Opportunity Gap, that is the difference between max and min weighted macro TPR across groups, shows that 3SD is the only model with equal performance across all groups (Figure 7A; see Methods). Similar results are obtained considering Predictive Equality Gap, that is the difference between max and min weighted macro FPR across groups (Figure 7B; see Methods).

**Figure 7.**
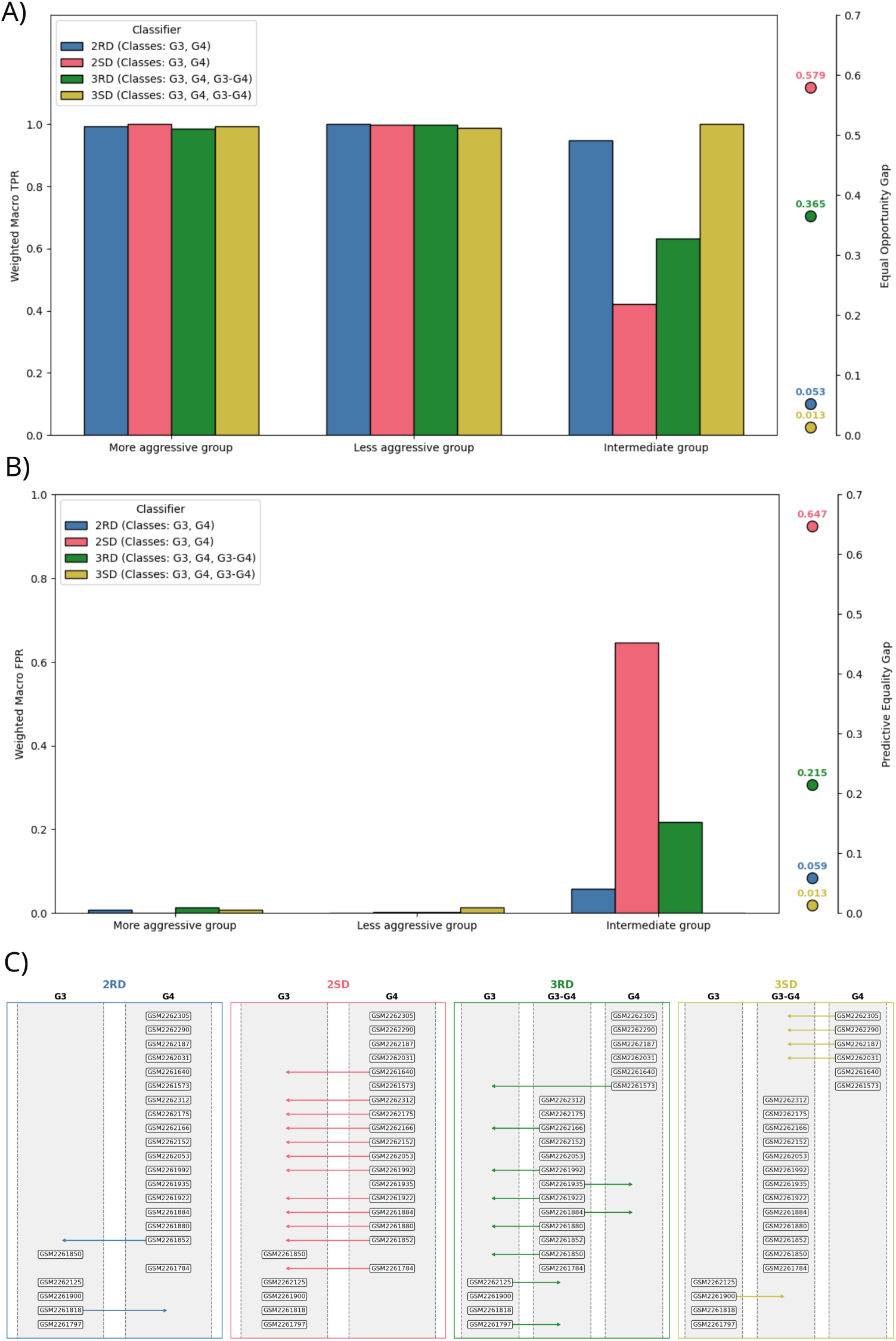
Fairness analyses. A) Equal Opportunity across groups. The barplot shows the weighted macro True Positive Rate (TPR) of two binary classifiers (2RD, 2SD) and two multi-class classifiers (3RD, 3SD) within three defined patient groups (More aggressive group, Less aggressive group, Intermediate group). Details on training and testing procedures are provided in the Methods section. The values of the Equal Opportunity Gap of each classifier are displayed on the right end side. **B) Predictive Equality across groups.** The barplot shows the weighted macro False Positive Rate (FPR) of two binary classifiers (2RD, 2SD) and two multi-class classifiers (3RD, 3SD) within three defined patient groups (More aggressive group, Less aggressive group, Intermediate group). Details on training and testing procedures are provided in the Methods section. The values of the Predictive Equality Gap of each classifier are displayed on the right end side. **C) Visualization of misclassified patients across classifiers.** Each panel corresponds to a different classifier (2RD, 2SD, 3RD, 3SD), with patients shown at their true label positions and arrows pointing to their predicted labels. Background columns represent the possible class labels in each context. Only patients with at least one misclassification are included. Arrows are color-coded by classifier to highlight misclassification patterns.

### 2.7. Members of the Intermediate Group Are Systematically Misclassified by Other Classifiers

We observed that certain patients are consistently misclassified as false positives (FPs) across multiple classifiers, except for 3SD (Figure 7C; see Methods). Notably, all of these patients belong to the identified Intermediate group. Specifically, patient GSM2261852 was misclassified by both the 2RD and 2SD classifiers, while patients GSM2261880, GSM2261884, GSM2261992, GSM2261922, and GSM2262166 were misclassified by both the 2SD and 3RD classifiers.

To further investigate, we examined the misclassification profiles of each classifier individually. The 2RD classifier misclassified two patients: GSM2261818, labeled as G3 and classified as G4, and GSM2261852, labeled as G4 and classified as G3. The latter, a member of the Intermediate group, was also misclassified by the 2SD classifier. Importantly, 2SD, a binary classifier trained on synthetically balanced data, showed the highest false positive rate within the Intermediate group (Figure 7B), erroneously identifying G4 patients as G3, including GSM261784, GSM2261852, GSM2261880, GSM2261884, GSM2261992, GSM2261922, GSM2262053, GSM2262152, GSM2262166, GSM2262175, GSM2262312, and GSM2261640.

In the case of the 3RD classifier, a non-negligible FP rate persists (Figure 7B), likely attributable to class imbalance inherent in the real data. By contrast, the 3SD classifier, which employs synthetic class balancing across three classes, did not produce any FPs within the Intermediate group. Its only misclassifications involved one G3 patient (GSM2261900) and four G4 patients (GSM2262031, GSM2262187, GSM2262290, GSM2262305). These misclassifications highlight the inherent challenges in defining boundaries between tumor grades, underscoring the complexity of patient stratification and the need to reassess current criteria, as demonstrated by this study’s comparison of a standard binary labeling with a three-label labeling emerging from an unsupervised approach (clustering).

## 3. Discussion

Medulloblastoma (MB) is a rare and heterogeneous disease, with transcriptomic studies often constrained by sample sizes, typically encompassing fewer than 200 cases (1, 14, 15, 16). MB is classified into four molecular subtypes: SHH, WNT, Group 3 (G3), and Group 4 (G4) (50). However, due to the varying incidence rates of these subgroups, even larger datasets exhibit an uneven distribution, with particularly minimal representation of patients showing overlap between G3 and G4 transcriptomic features (13). These patients are of particular interest, as recent evidence suggests they may represent a distinct subgroup with unique molecular and clinical characteristics (10, 13, 14, 15), and prognosis. In particular, G3 is typically more aggressive than G4 (41), thus misclassifying G3-G4 patients could lead to inappropriate treatments (42, 43). To date, limited data availability has constrained robust analyses needed to validate this hypothesis, prompting authors to refrain from drawing definitive conclusions regarding the existence of a potential G3–G4 subgroup (10). Expanding the availability of high-quality, well-annotated datasets remains crucial for enabling subgroup-specific investigations and advancing our understanding of MB biology.

In this work, we have resorted to generative approaches to augment transcriptomic data of patients who potentially belong to a new subgroup, called G3-G4, that would be otherwise mislabeled in traditional classifications. To address this challenge, we developed a VAE model, trained on the largest available transcriptomic dataset of MB patients, designed to capture a biologically meaningful low-dimensional representation of the data and generate synthetic patient profiles, facilitating the refinement of the disease’s molecular classification.

An unsupervised clustering approach, developed for this work, revealed a subset of patients that could not be definitively assigned to either G3 or G4. To augment this putative G3-G4 subgroup and at the same time balance out all the other subgroups, we generated high quality synthetic data using our VAE. When comparing the classification performance of synthetic data balanced using two labels (G3 and G4) versus three labels (G3, G3-G4, G4) high separability is maintained with a finer stratification. This is likely because the putative G3-G4 group contains few data points, which, even if potentially mislabeled as G3 or G4, have a minimal impact on the two-class performance.

Differential expression analysis among the G3, G4, and G3-G4 subgroups revealed genes previously implicated in MB pathology (45) (KMT2C, MYC, SNCAIP, SYNCRIP, TP53). One of the worst clinical outcomes in MB is found in MYC-amplified Group 3 patients (1, 41). Precisely, we found MYC to be one of the most important genes to be differentially expressed in the newly-found G3-G4 subgroup, potentially associated with a poorer prognosis than G4. Therefore, while further research is needed, our findings may prove valuable in preventing future mislabeling of patients.

To ensure synthetic data reliability and enhance its applicability, it is crucial to validate synthetic data not only through domain expert review, statistical consistency checks, and model performance and fairness testing, but also by comparing it with external, independent evidence. We performed an external validation comparing our findings with those reported in our earlier study using a different multi-omics dataset and a different methodology based on mul-tilayer networks (14). Strikingly, both approaches support the existence of a potential new subgroup G3-G4 with overlapping relevant genes (CDKN2C, IMPDH1, SLC35A4, ST8SIA2, ST8SIA3). Given the sensitivity of clinical practice, comparing results from generative and non-generative approaches is critical to ensure that the synthetic data are biologically relevant and reliable. Importantly, our Boolean-network analysis provides an orthogonal, mechanistic validation of the subgroup structure inferred by the VAE. While the representation-learning and synthetic-data framework operates at the transcriptomic level, the Boolean simulations show that G3, G3-G4 and G4 samples occupy distinct places in signaling space, with Group 3 dominated by MYC-driven proliferation/cell-cycle activation and relatively weaker PI3K-AKT-mTOR and motility signaling, and Group 4 displaying the converse pattern, with enhanced PI3K-AKT-mTOR and Rho-GTPase/PKC-centred cytoskeletal and motility pathways. The intermediate G3-G4 subgroup, which is essential for fairness-aware classification, consistently exhibits pathway activities that lie between these extremes, rather than collapsing onto either Group 3 or Group 4 signaling, and is concordant with the single-cell gene-regulatory network view of Group 3/4 tumors as a continuum along photoreceptor/MYC versus precursor/unipolar brush cell-like axes rather than strictly discrete subgroups (56). This multi-scale convergence from latent transcriptomic representations and synthetic cohorts to pathway activation patterns strengthens the case that G3-G4 represents a biologically meaningful state along the Group 3/4 continuum, with potential implications for both risk stratification and targeted therapeutic strategies.

Importantly, we conducted a thorough fairness analysis across all classifiers. Among all models, the balanced synthetic three-class classifier, explicitly accounting for the intermediate G3–G4 subgroup, demonstrated the highest fairness, achieving the most equitable error rates across patient groups. This work extends the principles of responsible AI beyond traditional social discrimination categories such as sex, gender, or race, addressing critical clinical concerns like treatment discrimination. Indeed, misclassification of patients within the intermediate G3–G4 group can have catastrophic consequences, given that G3 tumors typically require more aggressive treatment than G4 (64). In other words, patients in G3-G4 may not be receiving optimal care if their true tumor aggressiveness is not accurately reflected by their assigned labels leading to dangerous misclassifications, underscoring the urgent need for more precise stratification methods. These findings underscore the urgent need to integrate fairness assessments alongside predictive performance in clinical AI models, as advocated by initiatives such as FUTURE-AI (65).

A crucial aspect of applying machine learning models to medical problems is the need for explainability, especially when modeling molecular data with deep neural networks. Our explainability pipeline allowed us to identify the most important genes for subgroup classification by leveraging the latent variables learned during data synthesis. This is an important contribution given the notoriously challenging task of interpreting latent representations of molecular data compared to more directly human-readable data types, such as images or text.

## 4. Methods

### 4.1. Data Preprocessing

RNA microarray data from primary medulloblastoma patients was obtained from GEO under accession code GSE85217. The dataset contains the expression values of 21,641 genes in 763 patients that have been preprocessed and normalized. Genes were filtered out according to two criteria: genes with zero expression in at least 20% of the patients and genes whose expression mean and variance were below 0.5. Furthermore, outlier genes were identified and removed using the Mahalanobis distance (66). After preprocessing, the final dataset used for training and testing the VAE model contains the expression values of 14,349 genes in 763 patients.

### 4.2. Unsupervised Discovery of the G3-G4 Subgroup

To explore the possible existence of an intermediate group between G3 and G4, we attempted to replicate the procedure described in (10), for which no code is provided in the corresponding publication. This procedure consists in a consensus clustering of the top 10,000 most variable genes, trying to identify first two and then three groups. Patients who switched clusters between the two analyses were labeled as the G3-G4 subgroup. This procedure has been applied to the original GEO dataset and the preprocessed data, which removed lowly expressed genes, giving inconsistent results when compared to (10) (Figure S2A,B), and leading us to discard it from the analysis and develop a new methodology to identify the intermediate group.

The new methodology consists in a k-Nearest Neighbors Graph (k-NNG) followed by an agglomerative clustering to discover the optimal number of neighbors and clusters in the data from G3 and G4. Once the optimal number of neighbors and clusters is found, a bootstrapping approach is applied. This consisted of sampling without replacement 100 patients from each of the two groups, and applying the k-NNG and clustering to the sampled data. The process was repeated 1,000 times, counting the number of times each patient was assigned to a cluster other than its original one. Patients were considered G3-G4 when they were assigned to the other cluster more times than a predetermined cutoff value.

The cutoff value was defined as the expected value of a patient being assigned to the other cluster, given the number of patients in the two subgroups. A coin-toss probability was assumed to compare results to a purely random assignment. Hence, the probability of a patient being assigned to the other cluster was calculated as the ratio of the chosen number of patients times a 50% probability of being assigned to the other cluster. That is,

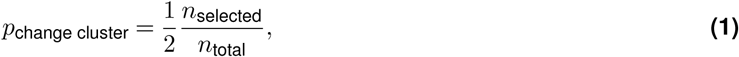

where *n_selected_* is the number of patients selected in each bootstrapping event (100 patients) and *n_total_* is the total number of patients in each subgroup: 144 in Group 3 and 326 in Group 4. With this probability, the expected value or cutoff for a patient being assigned to the other cluster was calculated as

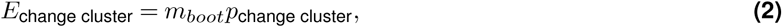

where *m_boot_* is the number of bootstrapping events (1,000). The resulting cutoff value is then 347.22 for G3 and 153.37 for G4.

Patients that changed clusters more times than the cutoff value were considered as G3-G4 patients.

### 4.3. VAE model training

We designed a VAE (21), an unsupervised method, with two hidden layers in both the encoder and decoder (model architecture shown in Figure S1). The encoder takes in as input dimension the number of genes, processes the data through a first hidden layer, and a second hidden layer estimates the mean *µ* and standard deviation *σ* of each dimension in the latent space. The reparametrization trick (21) was used to sample from one normal distribution *N* (*µ, σ*) per latent space dimension. The decoder then takes the sampled latent space, processes it in a complementary manner to that of the encoder, producing outputs in the original RNA microarray data space.

The preprocessed data was split into training and testing sets using an 80/20% ratio, stratified by patient subgroup, and normalized using the MinMaxScaler function from the scikit-learn library (67). The VAE was trained with the Adam optimizer, using a learning rate of 10*^−^*^4^, a batch size of 8, and a linear *β*-cycle-annealing schedule (68) for the KL divergence loss. The training objective is to minimize the negative Evidence Lower Bound (ELBO), which takes the form

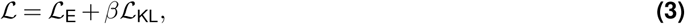

where L_E_ is the reconstruction loss, calculated as the Mean Squared Error (MSE), and L_KL_ is the non-negative KL divergence loss. The KL divergence loss was calculated between the approximate posterior and prior distributions,

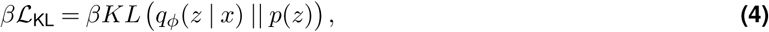

where *q_ϕ_*(*z* | *x*) is the approximate posterior distribution of the latent space *z* given the input *x*, parameterized by the encoder network weights *ϕ*, and *p*(*z*) is the prior distribution, set to a standard normal N(0*, I*).

The schedule consists of three cycles of 200 epochs each. Within each cycle, the *β* parameter increases linearly from 0 to 1 for the first 100 epochs and remains constant for the next 100.

### 4.4. Unsupervised Synthetic Data Generation

After training the VAE model, synthetic patients were generated in the latent space by sampling real, encoded patients and adding standard normal noise to them. Noise was scaled in the latent space with the MinMaxScaler function from the scikit-learn library (67) to match the range of the sampled patients. The scaled noise was adjusted by a noise-ratio factor and added to the sampled patients, creating a new set of patients in the latent space. The decoder then reconstructed these encoded, synthetic patients into the RNA microarray data space.

Once the VAE had been optimized and trained, it generated samples with a reconstruction loss that was unsatisfactory for our purposes: the most optimal architecture showed a reconstruction of 26.8 as the average of the last 20 epochs in the test loss (Figure S5A, C, and Supplementary Table S1). The implication of a moderate loss is that decoded samples from the latent space may exhibit slight distributional shifts compared to the real, input data. Newly generated synthetic data must achieve extreme distributional fidelity. In order to bridge a potential domain gap, we implemented a post-processing, unsupervised neural network with four hidden layers, trained on the VAE-decoded data to minimize the reconstruction error between the real and decoded data. During training and testing, we kept the same patient split as in the VAE model. The neural network was trained using the Adam optimizer with a learning rate of 10*^−^*^4^, a batch size of 8, and 1,000 epochs. The most optimal number of neurons for each hidden layer was obtained using Optuna (69) and were set to 2,981; 180; 3,699; and 14,349 neurons (the last one coincides with the number of genes after preprocessing). This additional step yielded reconstructed data that closely resembled the real data, lowering the reconstruction error from 26.8 to 0.04 on average for the last 20 test epochs (Supplementary Figure S6).

Our procedure is a two-step solution to train deep learning models, the style of which has been used in the literature (70, 71, 72, 73). The post-processing step acts solely on the reconstruction error of the VAE, leaving the KL-divergence intact, acting as a synthetic-domain alignment mechanism. The ‘reconstructed’ space was subsequently used to analyze synthetic data because of its resemblance to the original data space and because it retains the separability between the subgroups and the overlap between G3 and G4, as shown in the subgroup classification (Figure 2F) and reflected in the UMAP (Figure 2C). Moreover, the generated synthetic data overlaps with the real data in the UMAP (Supplementary Figure S10).

### 4.5. Differential Expression Analysis

To determine which genes were differentially expressed, two consecutive statistical tests were performed on the three subgroups of interest: G3, G4, and G3-G4. A Kruskal-Wallis test was used to determine if the expression of each gene was significantly different between the three medulloblastoma subgroups. Among the significantly different genes, a post-hoc Dunn’s test with Bonferroni correction was performed to identify in which subgroups genes were different from each other. The genes that were significantly different between all three subgroups were considered as differentially expressed. A p-value threshold of 0.01 was used for both tests.

Given the p-value depends on the sample size, we also calculated the effect size (Cohen’s d) on the genes identified as differentially expressed. We used the *pingouin* python package (74) to calculate the effect size and its 95% confidence interval. We followed general guidelines to determine the importance of the effect size (75, 76) as small (0.2 *< d* ≤ 0.5), medium (0.5 *< d* ≤ 0.8), and large (*d >* 0.8).

### 4.6. VAE Explainability Pipeline

To interpret the trained VAE, we calculated the SHapley Additive exPlanations (SHAP) values (32) for each input gene. This was done after training the VAE model, meaning we are interpreting the trained VAE externally. The explainability of the VAE model was performed in two steps: tracing the importance of latent variables to the medulloblastoma subgroups, and tracing the importance of input genes to the latent space.

First, a supervised classification of the four medulloblastoma subgroups (SHH, WNT, G3, and G4) on the latent space was performed with XGBoost (34) after hyperparameter optimization with Optuna (69). The classification returned a trained tree model that was explained using TreeExplainer (77) to map the importance of the latent space variables to each of the subgroups. Since we are interested in the overall importance, we studied the absolute value of the SHAP values. Given that SHAP’s TreeExplainer studies the importance of single predictions, the resulting SHAP values were averaged for each prediction and summed over all subgroups. The process was repeated 100 times to account for stochasticity in the classification and hyperparameter optimization. Hence, the obtained SHAP values were also summed over all repetitions, ending with one SHAP value per latent space variable. The top most important latent variables were selected among all latent variables as those whose absolute SHAP value is above the average (Supplementary Figure S11A). Only these most-important variables were considered for the next step.

Second, to assess the relationship between input genes and latent space variables, DeepExplainer (32) was used to map the importance of each gene. Here, the objective was to understand how the latent space variables were related to the gene expression. To do so, the DeepExplainer was executed considering all input genes and the encoder part of the VAE model, which is stochastic, given the step of sampling Gaussian distributions in the latent space. Because of this stochasticity, the DeepExplainer algorithm was repeated 100 times to account for the randomness in the VAE model. All genes kept after preprocessing were considered in the SHAP analysis. As in the previous case, the SHAP values, in absolute value, were averaged over all patients and summed over latent space variables and repetitions (Supplementary Figure S11B).

### 4.7. Enrichment Analysis

The enrichment analysis was performed using the gprofiler2 (55) package in R. The analysis aimed to identify significantly enriched biological terms and pathways among genes important for medulloblastoma subgroup classifi-cation, as determined from our explainability pipeline (Results 2.4 and Method 4.6). The enrichment was performed on annotated databases, namely Gene Ontology (GO) (78), Reactome (REAC) (79), Kyoto Encyclopedia of Genes and Genomes (KEGG) (80), and WikiPathways (WP) (81). The results of the enrichment analysis (Supplementary Dataset 4) include terms that are significant in humans, with a p-value below 0.05, corrected for with false discovery rate, considering annotated genes. The log-transformed p-values (−*log*_10_(*p*-*value*)) were calculated for visualization purposes. Supplementary Figure S12 shows the top 20 most significant terms from each of the annotated databases used.

### 4.8. Fairness Assessment

To evaluate the fairness of a classifier, we employed the Equalized Odds criterion (57), which requires equality of both True Positive Rates (TPR) and False Positive Rates (FPR) across sensitive groups. For a binary classification task, Equalized Odds is formally defined as:

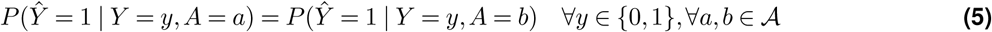

where *Ŷ* denotes the predicted label, *Y* the true label, and *A* the sensitive attribute with groups *a, b* ∈ A. This implies equal True Positive Rate (TPR) and False Positive Rate (FPR) across groups:

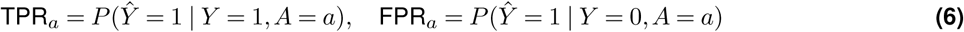

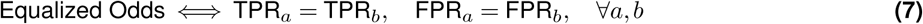

For multi-class classification with classes *k* ∈ {1*,…, K*}, we generalized Equalized Odds by computing *weighted macro-average* TPR and FPR per group, thus adjusting for class imbalances (62):

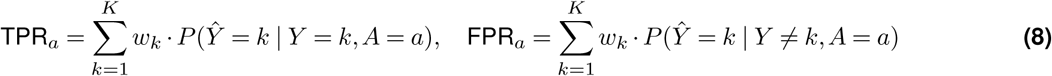

where weights 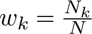 represent class support, with *N_k_* the number of instances of class *k*, and 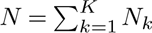.

Equality of these weighted metrics across groups satisfies multi-class Equalized Odds:

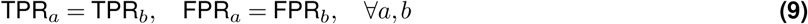

In this study, groups are defined based on the unsupervised approach that identified three clusters among all patients in Cavalli et al. (10): a newly identified cluster of patients that we labeled G3–G4 (Intermediate group), and the other clusters, consisting of the remaining G3-labeled individuals (More Aggressive group) and G4-labeled individuals (Less Aggressive group), respectively. The binary classifiers were trained using the labels G3 and G4 as defined by Cavalli et al. for the entire dataset as true labels, while the multi-class classifiers were trained using these same labels along with an additional G3–G4 label, which was assigned to individuals originally labeled G3 or G4 by Cavalli et al. but identified within the Intermediate group.

All classifiers were trained on real datasets and synthetic datasets and evaluated using Leave-One-Out CrossValidation (LOOCV) (63) to ensure unbiased performance estimates. For each fold *i* (where each instance is held out once):

1. Train the classifier on *N* − 1 instances, optimizing hyperparameters using internal validation within the training fold.
2. The trained model was evaluated on the held-out instance (i.e., each patient from the original dataset). For classifiers trained on synthetic data, the training set did not include any real patient instances.
3. Aggregate predictions over all folds for fairness assessment.

This protocol prevents data leakage and optimizes hyperparameters independently per fold, ensuring a fair comparison across models.

### 4.9. Boolean model construction

The focus of the model was to study the differences and similarities of medulloblastoma subgroups G3 and G4. We built a network with the genes and proteins identified as relevant in Forget et al. (15). Figure 5 of said paper focused on PI3K-AKT, mTOR, MAPK and RHO signaling pathways and shows G4/G3 ratios for mRNA, protein, and phosphorylation levels across ERBB1-4 receptors, G protein coupled receptors (GPCRs), and downstream effectors. These proteins provided the seed to construct our Boolean model, where protein activity is abstracted as active or inactive and this activity depends on the interactions between the genes, mRNAs, and proteins.

To connect this list of genes and proteins, we used NeKo (82), a tool for automatic network construction using the interactions from the SIGNOR database (83). Then, we wanted the model’s outputs to be cancer-related phenotypes (such as Apoptosis, Proliferation, and Metastasis). For this, we expanded the network by allowing NeKo to include other additional nodes that would connect the terminal nodes (for instance CFL1, CREB1 and mTORC1) to these phenotypes. We refined the Boolean network to have biologically proven interactions, while simplifying as much the number of these, and trying to keep the model computationally tractable within our means.

We further refined the model manually by consolidating protein complexes (mTORC1, mTORC2, PI3K, TSC) by merging individual components into functional units while preserving their regulatory logic; eliminating redundant protein family nodes (LATS1_2, AKT family) in favor of more granular individual isoforms with stricter AND activa-tion rules; and removing poorly integrated terminal nodes (Cell_cycle_progress, CDKN1A, RAC2, CDC42EP4) that lacked meaningful connections to the core phenotypes (Apoptosis, Proliferation, and Metastasis). These changes were backed up by SIGNOR or OmniPath databases, particularly for cytoskeletal regulation and migration pathways, while ensuring species-specificity to Homo sapiens. Logical formulas were build for each node following the parsimonious rule of adding the activators with AND gates and inhibitors with OR gates. Nevertheless, some nodes had their logical formulas adapted so they were sensitive to inputs, like PRKCA, ARHGEF1, RHOA, AKT isoforms (see Supplemental File X for details). For instance, RHOA was edited as its permissive inhibitors logic combined with dense positive feedbacks loops caused constant activation of the outputs regardless of the inputs’ initial states. The resulting network consisted of 111 nodes (including 8 inputs and 3 outputs) and 344 edges was then converted into .BND and .CFG files with bioLQM (84) to be used in MaBoSS to simulate the Boolean model probabilistically.

### 4.10. Boolean model personalization

We personalized the model by adapting the PROFILE pipeline (85) to use microarray data. This methodology uses omics data to constrain the probabilities of MaBoSS, which uses a continuous-time Markov process on the state transition graph of the Boolean model (see (86) for details). Most genes had a one-to-one relation with a node in the model, for the ones that had not, like complexes, we selected the minimal component for each complex and assigned it to the respective complex. Genes were classified in the Unimodal category using the new compute_criteria_microarray function. Normalized expression values were then used as initial conditions and as transformation rates by using the following formula:

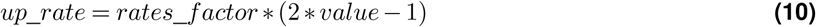

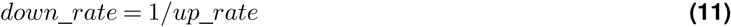

Where rates_factor corresponds to the argument rf which in this case we set at 100. The initial states of the nodes were set at random in the wild-type model and after the personalization for each patient, some input nodes were modified according to their normalized transformed expression value, with the following formula:

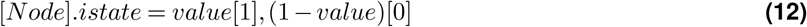

### 4.11. Model simulation and analysis of results

Finally, we simulated the models using MaBoSS, version 2.6, (86) and pymaboss interface (84). We set use_psyrandgen = FALSE and seed_pseudorandom = 100 to enable fast, reproducible pseudo-random trajectories; max_time = 60 as trajectories stabilize well before this point; time_tick = 0.1 to improve temporal resolution within the shortened window; and statdist_traj_count = 100 to robustly quantify trajectory distributions. All other parameters remained at the default values. We simulated the 763 patients from the microarray data from Cavalli et al. (10) and the 600 synthetic patients generated by this work using the Garnatxa computing cluster at I2SysBio.

For pathway-level analyses, we first defined biologically coherent node sets representing major signaling modules (for example, Apoptosis, Proliferation-cell cycle, PI3K-AKT-mTOR, RAS-MAPK, cytoskeleton/actin dynamics, Invasion-motility, Hippo-YAP, receptor tyrosine kinase signaling).

Then, all statistical analyses and visualizations were performed in R (v4.5.3). For each sample and pathway, we used the node-level activation probabilities from the MaBoSS simulations and computed a pathway activation score as the arithmetic mean of the stationary activation probabilities of all nodes assigned to that pathway. These persample pathway scores were merged with the group labels and used as input for all downstream statistical testing and plotting. To assess the differences in pathway activation between group classes, we applied non-parametric tests. For each pathway, we first performed a Kruskal-Wallis test across all six group categories to test for any global differences in activation. When this test was significant, we conducted post hoc pairwise Mann-Whitney-Wilcoxon rank-sum tests between groups (for example, G3 vs G4, G3 vs synthetic G4, synthetic G3 vs synthetic G4, etc.), with Benjamini-Hochberg adjustment of p-values to control the false discovery rate across comparisons. In parallel, we quantified effect sizes for key contrasts using Cohen’s d, calculated as the difference in group means divided by the pooled standard deviation, to complement p-values with an estimate of the magnitude of between-group separation on the scale. All tests were two-sided, and both adjusted p-values and effect sizes were reported for the interpretation of biologically relevant differences.

For Figure 6, we generated a clustered heatmap of the mean pathway activation by group. To this end, we first summarized the per-sample pathway scores by computing the mean value of each pathway within each group class, yielding a pathway-by-group matrix. This matrix was visualized with the ComplexHeatmap package using Euclidean distance and complete linkage hierarchical clustering on the pathway (row) and groups (column) dimensions. We scaled the mean values per pathway (row-wise z-scores) so that colors reflect relative increases or decreases within each pathway across groups rather than absolute activation levels. The resulting z-scores were color-mapped with a diverging blue-white-red palette (blue = low, red = high). Group labels and pathway names were added as annotations to facilitate their interpretation.

## 6. Data Availability

The dataset analyzed in this study is publicly available through the Gene Expression Omnibus (GEO) under the accession code GSE85217. A script to download the dataset is provided in our GitHub repository (see Code Availability)—an active internet connection is required to execute the script. The generated synthetic data and metadata are available to download at https://zenodo.org/records/14516824.

## 7. Code Availability

The code is available on https://github.com/gprolcastelo/synthetic_medulloblastoma.

## 9. Authors Contribution

G.P.C. and A.T.L. developed the core methodological concepts and technical implementations, and wrote and revised earlier and consolidated versions of the manuscript. G.P.C. and B.U. identified the datasets and handled the preprocessing of the microarray data. I.N.C. supervised the functional interpretation of the results. E.G.V and A.M. developed the Boolean model, its personalization, simulation, and analysis. A.V. and D.C. conceived the study and oversaw its execution. All authors reviewed and contributed to the final manuscript.

## Acknowledgements

The authors acknowledge the donors and families of the data used in this analysis. This work has been supported by the EU project EVENFLOW under Horizon Europe agreement No. 101070430. E.G.V and A.M. acknowledge funding from the Generalitat Valenciana’s CIDE GenT and ES GenT programmes under the project numbers CIDEXG/2023/22 and CIESGT/2025/02 and the technical help from Vincent Noël, from Institut Curie, to compile MaBoSS in the ARM64 architecture.

