## Supplementary Material for "Explainable Generative AI Uncovers a Molecular Continuum in Medulloblastoma with Implications for Rare Cancer Subtyping and Treatment Equity"

### Supplementary Information

#### Supplementary Text

##### *Model construction.*

**Initial node list** The model was constructed by first defining an initial list of seed nodes to be used as input to NeKo. NeKo connects seed nodes through a selected interaction database; in this study, the SIGNOR database **??** was used as the primary source of curated regulatory relationships. The initial node list was largely based on the network shown in Figure 5 of Forget et al. (2018) (1), with additional G protein coupled receptors (GPCRs) included when they were highly expressed in the non-WNT/non-SHH group described by Whittier et al. (2013) (2), provided that they were also present in SIGNOR.

The first NeKo-generated network contained disconnected nodes and nodes without upstream regulators, which limited its interpretability and prevented a coherent definition of inputs (ligands or receptors), intermediates, and outputs (phenotypes). To address this, intermediate nodes were introduced where necessary so that the resulting network could be organised into a connected signalling structure with explicit input, intermediate, and output layers. This step was important to avoid isolated nodes behaving as artificial inputs in the Boolean model.

A manual curation step was also performed to incorporate GIT1 as an upstream regulator of ARHGEF8, based on evidence curated in SIGNOR. This modification ensured that ARHGEF6 remained properly connected within the network and preserved the coherence of the downstream signalling architecture. In summary, the final input list combined nodes from Forget et al. (2018) (1), selected GPCRs from Whittier et al. (2013) (2), and the manually added regulator GIT1. All networks were exported as SIF files until we merged them together in a BNET file, used for its simulation.

**Intermediate node identification between GPCRs and ARHGEFs** To connect the GPCR layer to downstream effectors, NeKo was allowed to introduce intermediate nodes and found a single bridging node, GNA13, which links the GPCR set to the ARHGEF module from Forget et al. (2018) (1) composed of ARHGEF1, ARHGEF11, and ARHGEF12.

To maintain model interpretability and avoid excessive complexity, we restricted the network to this subset of nodes rather than allowing the inclusion of a large number of additional intermediates. This decision prevented the expansion of the model into an overly large and computationally-challenging network, facilitating analysis and simulation.

**Phenotype integration** Curated SIGNOR relationships between proteins and phenotypes were used to connect terminal signaling nodes to biologically relevant cancer phenotypes. Cancer-associated phenotypes were selected and linked to the network using NeKo, again permitting intermediate nodes when required to reach them. For each node subset, the resulting networks were merged into a single BNET file used for the simulations.

**Logical rule construction** Logical formulas governing the activation state of each node were systematically constructed from the curated regulatory interactions according to the canonical Boolean logic for regulatory networks. For each node, activating incoming edges were combined using AND logic, while inhibitory incoming edges were combined using OR logic.

This formulation reflects the biological principle that multiple positive signals are typically required for pathway activation, whereas any single inhibitory signal is sufficient to prevent node activity (3). Formally, for a node X with activating regulators A_1,A_2,. . . ,A_n and inhibitory regulators I_1,I_2,. . . ,I_m, the logical rule was defined as:

$$X = (A_1 \wedge A_2 \wedge ... \wedge A_n) \wedge \neg(I_1 \vee I_2 \vee ... \vee I_m)$$

When no activating regulators were present, the AND clause was omitted (reducing to $\neg(I_1 \vee ... \vee I_m)$). Similarly, when no inhibitory regulators were present, the inhibition term was omitted (reducing to $A_1 \wedge ... \wedge A_n$). Nodes lacking both activating and inhibitory inputs were assigned a default constitutive activation rule (X=1) unless biological evidence justified an alternative formulation.

This systematic AND-for-activation, OR-for-inhibition rule was applied uniformly across all nodes during network curation, ensuring consistent logic throughout the model while preserving the hierarchical and combinatorial nature of biological signaling. The resulting Boolean formulas were incorporated into the final BNET file used for MaBoSS simulations, providing a mechanistically interpretable representation of the regulatory architecture that balanced fidelity to curated interactions with computational tractability.

***Network curation.***

**Complexes and protein families** Several structural modifications were introduced before simulation to improve biological realism and reduce redundancy. MAPKAP1 was incorporated as an essential component of mTORC2 in line with Forget et al. (2018) (1). Although the explicit component-level representation of the mTOR complexes was later removed, MAPKAP1 remained conceptually part of mTORC2 during the personalisation step.

To simplify the network, the individual PIK3CA and PIK3R1 nodes were replaced by a single PI3K node representing the functional complex with a merged logical formula. Likewise, TSC1 and TSC2 were consolidated into a single TSC node. These changes reduced complexity while preserving the functional logic of the corresponding complexes. The LATS1_2 family node was removed because LATS1 and LATS2 were already represented individually, making the family-level node redundant. The AKT family node was also removed for the same reason, as AKT1, AKT2, and AKT3 were all explicitly represented.

**Biological process nodes** Nodes that were weakly integrated or functionally redundant were removed where appropriate. Cell_cycle_progress and CDKN1A were excluded because they formed a poorly connected submodule with limited relevance to the apoptosis, proliferation, and metastasis phenotypes analysed. Retaining such terminal or inhibitory-only nodes would have increased complexity without adding mechanistic value, and could have led to unrealistic Boolean behaviour if nodes lacking activators became constitutively active in the absence of inhibition.

Several process-level connections were added to improve biological coherence. Cell_migration was defined as an activator of Metastasis, capturing the principle that migration can promote metastatic potential. Neuron_migration was linked to Cell_migration, reflecting its specialised motility programme. Actin_cytoskeleton_regulation was also connected to Cell_migration, consistent with the role of cytoskeletal remodelling in motility. In addition, Protein_-synthesis was connected to Proliferation to represent the requirement for biosynthetic capacity during cell growth and division.

To enforce mutual exclusivity between survival and death phenotypes, Metastasis and Proliferation were modified to require the absence of Apoptosis. The Survival node was removed because it was redundant with Apoptosis in this model framework and did not contribute additional mechanistic information.

**F-actin assembly** The original logical formulation of F_actin_assembly was revised to better reflect actin dynamics. Rather than treating PAK1 as the sole activator opposed by cofilins, the node was redefined as F_actin_dynamics to capture the combined contribution of PAK1, CFL1, and CFL2 to actin remodelling. This change reflects the fact that filament turnover and polymerisation are jointly required for the formation of migratory structures such as lamellipodia and filopodia, rather than representing a purely unidirectional assembly process.

**Addition of regulators and removal of isolated nodes** Additional regulators were introduced where nodes would otherwise lack upstream control. PAK3 and PAK4 were assigned CDC42 as an upstream activator based on OmniPath evidence. ATF2 was linked to MAPK1 and MAPK3, and NFE2L2 was assigned upstream regulation by PRKCA, PRKCB, BRAF, and MYC, with PAK4 retained as an inhibitor. ROCK2 was defined as a RHOA target, and ARHGEF and the ROCK proteins were connected to Actin_cytoskeleton_reorganization according to SIGNOR annotations.

Further refinements included defining ARHGEF6 as an activator of Cell_migration and PRKCA as an inhibitor of Apoptosis. YES1 and EPHA2 were added as activators of WWTR1, and the ITCH–SPART–WWP1 regulatory chain was incorporated to control AMOTL2. This ensured that the YAP/TAZ-related module was embedded within the broader signalling context.

Nodes lacking positive regulators were systematically revised to prevent constitutive activation. Tubulin was connected to TTL through an “inhibitor-of-an-inhibitor” logic in which STMN inhibits TTL, thereby allowing Tubulin activation in a biologically interpretable manner. CDK1 was added upstream of STMN, and the upstream control of CDK1 was completed through CDC25B and PRKACA/RPS6KA1. Additional chains were introduced for FOXO through SIRT1, and FUS was linked to EGFR and mTORC2.

Finally, several nodes that lacked meaningful upstream integration and did not contribute to the phenotypes of interest were removed, including RAC2 and CDC42EP4. These pruning steps reduced unnecessary complexity without compromising the connectivity of apoptosis, proliferation, or metastasis-related modules.

**Logic refinement and species-specific curation** Several logical rules were refined to reduce constitutive activation and better reflect combinatorial regulation. PRKCA and ARHGEF1 were converted from permissive OR-based logic to more restrictive AND-based logic, thereby requiring coordinated inputs for activation. RHOA was also refined to represent combined inhibitory control by PRKACA and SRC in a more mechanistically consistent manner.

Interactions that were species-specific or not applicable to human biology were removed after checking the provenance of the NeKo-generated edges. In particular, PTEN-to-CREB1 inhibition and CREB1-driven proliferation were excluded because they were documented only in mouse rather than human systems. This curation step ensured that the final model remained restricted to human-supported regulatory relationships.

***Pathway-level analysis of the Boolean model results.***

**Data structure and preprocessing** The MaBoSS output matrix comprised one row per medulloblastoma sample and one column per Boolean-network node, with node values bounded between 0 and 1 and interpreted as stationary-state probabilities of node activation. Grouping2 was used as the primary biological class assignment variable throughout the comparative analyses. All remaining columns were treated as quantitative node-activation features.

Prior to statistical analysis, columns were partitioned into sample identifiers, class labels, and node-activation variables. No transformation was applied to the activation probabilities before pathway aggregation, because the original scale already represented model-derived probabilities and was directly comparable across samples within the same network framework.

**Pathway-level abstraction** To obtain pathway-level read-outs from node-level MaBoSS probabilities, biologically related nodes were grouped into predefined signalling and process modules. These included apoptosis, proliferation/cell cycle, PI3K-AKT-mTOR signalling, RAS-MAPK signalling, Hippo-YAP signalling, receptor tyrosine kinase signalling, cytoskeleton/actin dynamics, and migration/invasion.

For each sample, a pathway activation score was calculated as the arithmetic mean of the activation probabilities of the nodes assigned to that pathway (row-wise z-scores) (Figure 6). When an abstract functional node representing a higher-level process was available in the Boolean network (for example, Apoptosis or Proliferation), it was retained within the corresponding pathway definition together with mechanistically related upstream nodes, so that the pathway score reflected both the process read-out and its associated signalling context.

**Group-wise statistical testing** Pathway activation scores were compared across grouping classes at two levels. First, an omnibus test was used to assess whether each pathway differed globally across the six classes. Because pathway scores are bounded and may not satisfy strict normality assumptions, a non-parametric Kruskal-Wallis test was used for across-group comparison.

Second, all pairwise group comparisons were evaluated for each pathway. For each pair of groups, the difference in pathway means was calculated together with Cohen's d, which was used as the principal standardised effect size. Cohen's d was computed as the difference between group means divided by the pooled standard deviation, allowing effect magnitudes to be compared across pathways with different baseline means and variances.

To assess whether pairwise differences were likely to be non-random, Mann-Whitney-Wilcoxon tests were performed for each pathway-by-group-pair contrast (Figure S14). Statistical significance labels were recorded using conventional thresholds (* for $p < 0.05$, ** for $p < 0.01$, *** for $p < 0.001$), while interpretation prioritized concordance between statistical support and effect-size magnitude rather than p-values alone.

**Effect-size interpretation** Cohen's d was preferred over raw mean difference for visual comparison across pathways because it standardizes each between-group difference by the pooled within-group variability. This makes pathways with different intrinsic dynamic ranges and variances more directly comparable in a single summary figure. Raw mean differences were retained in the pairwise results table, but graphical summaries focused on Cohen's d because it better reflects the strength of biological separation between groups.

The sign of Cohen's d was interpreted relative to the ordering of groups in each pairwise comparison. A positive value indicated higher pathway activation in 'Group1' than in 'Group2', whereas a negative value indicated the opposite. In directional network visualisations, this sign was used to orient edges so that arrows consistently pointed from the group with higher pathway activation towards the group with lower pathway activation for the pathway under consideration.

**Heatmap of pairwise effect sizes** To summarize pathway-level differences compactly, a heatmap of pairwise Cohen's d values was generated. In this figure, rows corresponded to pathways and columns to pairwise group comparisons, with the cell value representing the standardized magnitude and direction of the pathway difference between the two groups. A diverging color scale centered at zero was used so that positive and negative effects were visually distinguished, and optional clustering could be applied to reveal shared patterns among pathways or group contrasts.

This heatmap was intended as the principal supplementary overview of pathway-level effect sizes. It allows readers to identify whether biologically coherent modules, such as proliferation or PI3K-AKT-mTOR signalling, consistently distinguish particular medulloblastoma classes, and whether synthetic groups recapitulate the same directional differences observed among patient-derived classes.

**Reproducibility and reporting** All statistical analyses and visualizations were generated from the MaBoSS simulation matrix and the derived pathway definitions using scripted workflows deposited in the project GitHub repository. The repository contains the full pathway mapping, statistical workflow, figure-generation scripts, and the exact parameters used for each visualization. The supplementary figures should therefore be interpreted together with the accompanying pairwise results table and pathway definition table, which together provide a complete record of the transformations from node-level probabilities to pathway-level comparative summaries.

**Principal component analysis of personalised model simulations** To further characterise the personalised models, principal component analysis (PCA) was performed on the MaBoSS simulations of the 1363 models. The first two principal components explained 14.8% and 10.5% of the total variance, respectively (Figure S15. In the global PCA embedding, PC2 recapitulated the expected continuum from Group 3 to Group 4 signalling states: G3 and synthetic G3 (s_G3) samples were displaced towards negative values of PC2, G4 and synthetic G4 (s_G4) towards positive values, and both the real intermediate class (G3-G4) and its synthetic counterpart (s_G3-G4) occupied positions between these extremes. Notably, the most contributing variables to the PC2 are SRC, CDC42 and PAK4 on the positive side and MYC, NFE2L2 and RHOA on the negative side. This pattern is consistent with the pathway-level analysis, in which G3 samples exhibit higher Proliferation signalling and lower PI3K-AKT-mTOR and motility programmes than G4 samples.

Unexpectedly, PC1 separated real G3-G4 from synthetic s_G3-G4 samples' centroids, despite overlapping along PC2 and sharing very similar mean pathway activation profiles. The other real and synthetic groups' centroids overlapped in both PC1 and PC2. Inspection of the underlying pathway scores confirmed that, for G3-G4 versus s_G3-G4, mean activation levels were closely matched across all pathways (Figure S14), and there were no extreme outliers that could trivially explain the PC1 displacement (Figure S16). Pairwise Wilcoxon rank-sum tests for each pathway and the G3-G4 versus s_G3-G4 comparison yielded no statistically significant differences in central tendency (all $p > 0.05$, Figure S14).

To investigate whether differences in variability rather than mean activation might underlie this real-synthetic separation along PC1, we compared the dispersion of pathway scores between G3-G4 and s_G3-G4 using the Fligner-Killeen test. Across pathways, only RTK signalling showed significant differences in variance between G3-G4 and s_G3-G4 (Fligner-Killeen $p = 0.014$), whereas all other pathways exhibited comparable dispersions. This indicates that PC1 is sensitive to subtle changes in the spread of RTK signalling activities between G3-G4 and s_G3-G4, rather than to large shifts in their mean activation.

Consistent with this interpretation, correlation analysis between PC1 scores and pathway activations within each class showed that, for G3-G4 and s_G3-G4, PC1 is strongly positively correlated with RTK signalling (Spearman $\rho = 0.696$ and $\rho = 0.394$ for G3-G4 and s_G3-G4, respectively), with PI3K-AKT-mTOR signalling (Spearman $\rho = 0.537$ and $\rho = 0.657$ for G3-G4 and s_G3-G4, respectively), and with Invasion/motility signalling (Spearman $\rho = 0.418$ and $\rho = 0.753$ for G3-G4 and s_G3-G4, respectively). Scatter plots of PC1 versus PC2 coloured by these pathways' scores illustrate that real and synthetic intermediate samples occupy partially segregated regions along PC1 that align with gradients in these two pathways, despite their similar mean values (Figure S17). Together, these observations suggest that the unexpected separation of G3-G4 and s_G3-G4 along PC1 predominantly reflects modest but systematic differences in the variability and joint distribution of PI3K-AKT-mTOR, Invasion/motility and

RTK signalling, rather than major shifts in pathway-level averages.

#### Supplementary Datasets

**Supplementary Dataset 1. Significantly differentially expressed genes limma.** limma-obtained differentially expressed genes in the synthetic G3-G4 subgroup compared to synthetic Groups 3 and 4, with an adjusted p-value lower than 0.01.

**Supplementary Dataset 2. Significantly differentially expressed genes with Kruskal-Wallis and Dunn's tests.** Differentially expressed genes in the synthetic G3-G4 subgroup compared to synthetic Groups 3 and 4, with a p-value lower than 0.01. Includes the effect size and its corresponding 95% confidence interval (CI).

**Supplementary Dataset 3. Important genes for subgroup classification.** Genes found to be important by our SHAP-based approach for the classification of the four original medulloblastoma subgroups (SHH, WNT, Group 3, Group 4) in the latent space. The importance was mapped from the subgroup to the latent variables, and then to the genes, as described in Methods.

**Supplementary Dataset 4. Enrichment analysis results.** This dataset includes the terms and pathways that were found to be significantly enriched. The analysis was performed with *gprofiler* (4) on the genes determined to be important for classification of the original medulloblastoma subgroups (see Methods).

**Supplementary Dataset 5. Boolean model files.** This dataset includes the BND and CFG files necessary to simulate the Boolean model build in this work (the generic and the 1363 personalised models) and a spreadsheet with the description of all the nodes, their source, their formula, and connections (see Methods, sections 4.9, 4.10, and 4.11).

**Supplementary Dataset 6. Boolean model statistical results files.** This dataset includes four tables: pathway_group_-means.csv, mean MaBoSS pathway activation probabilities across medulloblastoma groups; pathway_group_stats.csv, group-wise summary statistics of MaBoSS pathway activation scores; pathway_KruskalWallis.csv, global Kruskal-Wallis tests for differences in pathway activation between medulloblastoma groups, with Kruskal-Wallis H statistic, p-value and significance for each pathway; and pathway_pairwise.csv, pairwise group comparisons of pathway activation: mean differences, Cohen's d effect sizes and Mann-Whitney p-values with significance codes.

#### Supplementary Tables

| | 8 | 16 | 32 | **64** | 128 | 256 | 512 |
|---|---|---|---|---|---|---|---|
| 256 | 57.482651 | 46.329087 | 37.176923 | 35.749503 | 34.255623 | 45.636683 | 46.860007 |
| 512 | 40.576170 | 31.088746 | 29.134848 | 28.046977 | 27.854216 | 28.776864 | 34.039303 |
| **1024** | 35.857947 | 27.669830 | 26.915019 | **26.803298** | 27.279420 | 29.683946 | 42.642558 |
| 2048 | 35.924637 | 28.437002 | 27.539900 | 27.186680 | 28.765509 | 36.696050 | 60.568192 |
| 4096 | 36.779536 | 29.666869 | 28.980013 | 30.045203 | 30.417230 | 51.178306 | 81.946701 |

**Supplementary Table S1. VAE testing errors for different architectures.** The table shows the values for the mean of the last 20 epochs of the reconstruction test loss of the trained VAE across different model architectures. The table reflects the numbers used for displaying the corresponding heatmap in Sup. Fig. S5A. Columns represent the number of latent variables and rows represent the hidden layer dimension of the encoder and decoder parts of the VAE. The model with the lowest reconstruction loss is highlighted in bold.

| Original \ Clustering | Group 3 | Group 4 | G3-G4 | SHH | WNT | All |
|---|---|---|---|---|---|---|
| Group 3 | 142 | 0 | 2 | 0 | 0 | 144 |
| Group 4 | 0 | 309 | 17 | 0 | 0 | 326 |
| SHH | 0 | 0 | 0 | 223 | 0 | 223 |
| WNT | 0 | 0 | 0 | 0 | 70 | 70 |
| All | 142 | 309 | 19 | 223 | 70 | 763 |

**Supplementary Table S2. Contingency table between original and subgroups found with our bootstrapping method.** The table shows the number of patients in each subgroup, as determined by our bootstrapping method for clustering (Methods).

| WNT | SHH | Group 3 | Group 4 |
|---|---|---|---|
| CTNNB1 | TERT | **MYC** | PRDM6 |
| DDX3X | DDX3X | SMARCA4 | **SNCAIP** |
| SMARCA4 | KMT2D | KBTBD4 | KDM6A |
| CSNK2B | CREBBP | CTDNEP1 | ZMYM3 |
| **TP53** | TCF4 | KMT2D | **KMT2C** |
| KMT2D | PTEN | MYCN | KBTBD4 |
| PIK3CA | **KMT2C** | OTX2 | MYCN |
| BAI3 | FBXW7 | | OTX2 |
| EPHA7 | GSE1 | | CDK6 |
| ARID1A | BCOR | | GFI1 |
| ARID2 | PRKAR1A | | GFI1B |
| **SYNCRIP** | IDH1 | | |
| ATM | | | |

**Supplementary Table S3. Commonly mutated genes in each medulloblastoma subgroup.** This table lists the most commonly mutated genes for each medulloblastoma subgroup, as reported in (5). Genes we found to be significantly differentially expressed in the G3-G4 subgroup are highlighted in bold.

| Noise Ratio | Subgroup | Original | Reconstruct |
|---|---|---|---|
| 0.1 | SHH | 0.072637 | 0.071575 |
| | WNT | 0.083996 | 0.072938 |
| | Group 3 | 0.086572 | 0.073282 |
| | Group 4 | 0.070892 | 0.059620 |
| 0.2 | SHH | 0.223430 | 0.075184 |
| | WNT | 0.232439 | 0.075374 |
| | Group 3 | 0.235642 | 0.075274 |
| | Group 4 | 0.220979 | 0.061987 |
| 0.3 | SHH | 0.475756 | 0.078708 |
| | WNT | 0.482756 | 0.080643 |
| | Group 3 | 0.485179 | 0.078530 |
| | Group 4 | 0.472970 | 0.071629 |
| 0.5 | SHH | 1.143278 | 0.095206 |
| | WNT | 1.146574 | 0.093596 |
| | Group 3 | 1.153027 | 0.098370 |
| | Group 4 | 1.143232 | 0.112525 |

**Supplementary Table S4. Distance between real and synthetic subgroups.** This table shows the average Wasserstein distance between each of the real subgroups and their synthetically generated counterparts. In the case of column “Original", noise with varying ratios is added directly to the real data. Low levels return synthetic subgroups that are relatively close to the real subgroups. As the noise ratio increases, so does the distance between the real and synthetic subgroup. In the “Reconstruct" column, noise with varying ratios is added to the latent space, then decoded with the VAE and postprocessed with the reconstruction network (Methods). Even as the noise ratio increases, the distance between the real and synthetic subgroup stay similar.

#### Supplementary Figures

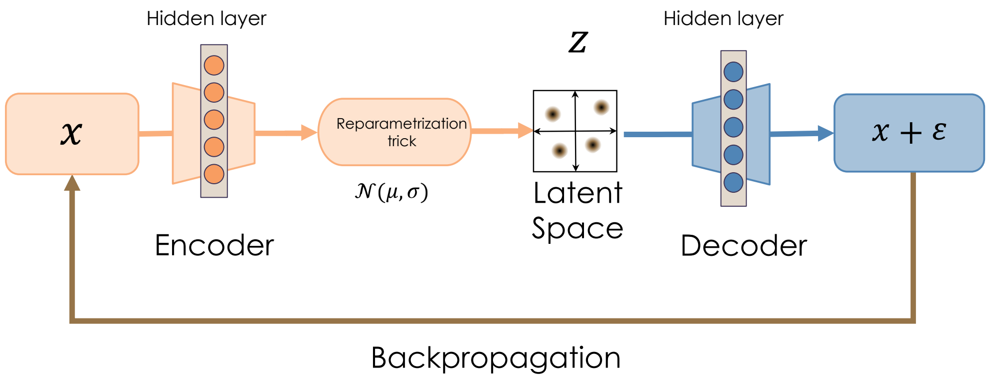


**Supplementary Figure S1. Diagram of a Variational Autoencoder (VAE).** $x$ represents the input transcriptomics data, $z$ denotes the latent space embedding of $x$. The reparametrization trick yields a normalized distribution $N(\mu, \sigma)$, where $\mu$ is the mean and $\sigma$ the standard deviation of the distribution for each latent space dimension. $\varepsilon$ is the training error, i.e., the difference between the real and reconstructed data. The VAE does not learn from the subgroups' labels, but they were used to stratify the input data x.

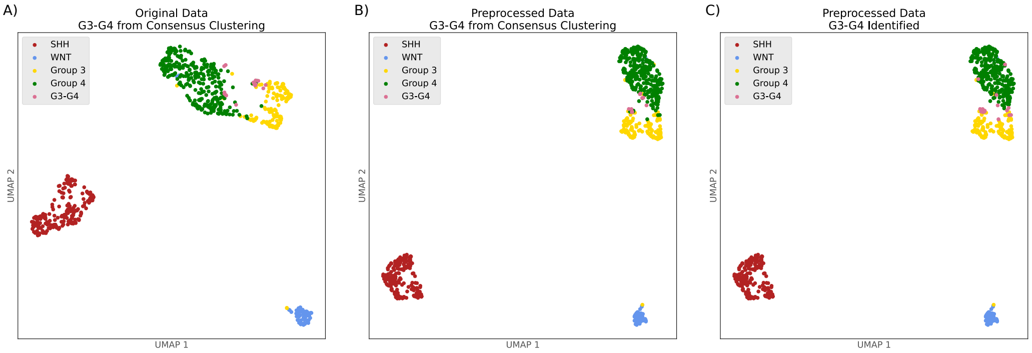


**Supplementary Figure S2. UMAP of identified G3-G4 subgroup with A),B) consensus clustering and C) our methodology.** The G3-G4 group was identified within Groups 3 and 4 with consensus clustering following the methodology described by Cavalli (6) for A) the original data from Cavalli, finding 15 patients at G3-G4, and B) the preprocessed data, finding 10 patients at G3-G4. C) The G3-G4 group was identified with the bootstrapping methodology we have developed (Methods), finding 19 patients at G3-G4. Meanwhile, the publication we obtained the data from (6) had determined that 14 patients were G3-G4, but did not provide the list of patients.

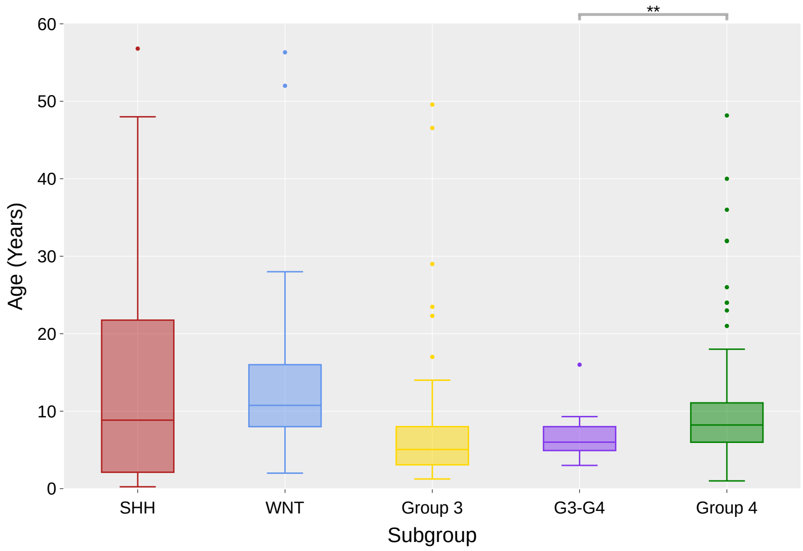


**Supplementary Figure S3. Age-at-diagnosis distribution between different subgroups, including the newly-identified G3-G4.** The figure shows the distribution in years of the patients age at diagnosis. A two-tailed Mann-Whitney U test was performed comparing the distributions of G3-G4 with Group 3 and, separately, with Group 4, taking both sexes into account. A significant difference was found with respect to Group 4 (U = 3664.5, p = 0.009, Cohen's d = 0.496), but not Group 3 (U = 1003.5, p = 0.249, r = 0.039). Asterisks indicate the significance level of the significant comparison between G3 and G3-G4 ($**p < 0.01$). Age data extracted from (6).

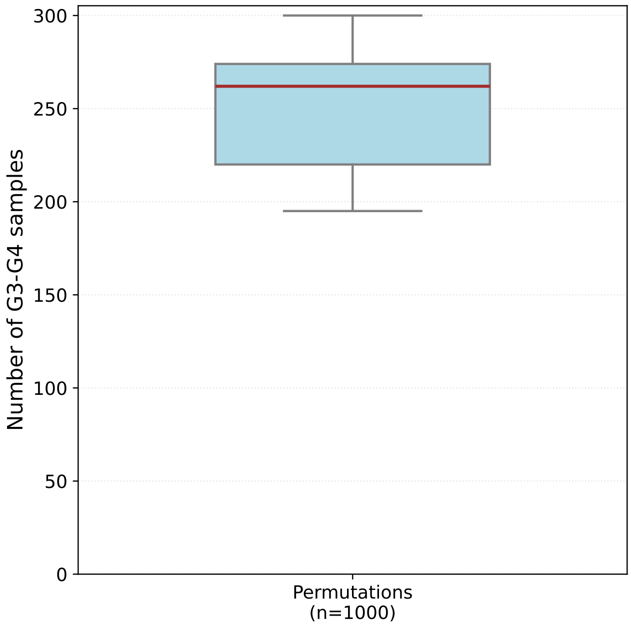


**Supplementary Figure S4. G3 and G4 label randomization experiment.** Distribution of the number of patient samples found to be intermediate between G3 and G4 by performing a random shuffling of the original labels. The experiment was repeated 1,000 times. The red, bold line indicates the median number of samples found to be intermediate by the knn with bootstrapping experiment, which, with the original labels, found 19 samples. The median of the distribution is 262 samples, and the standard deviation is 34 samples.

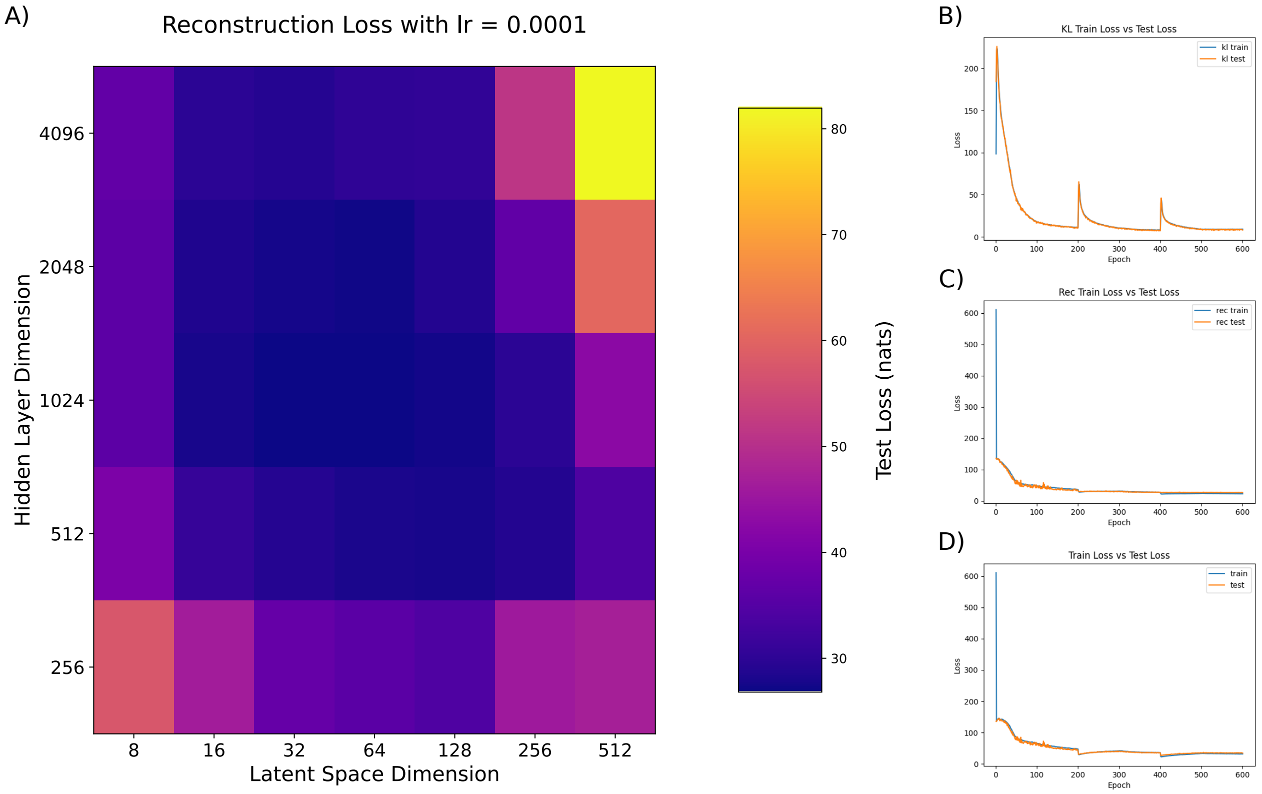


**Supplementary Figure S5. VAE performances.** A) Heatmap of Reconstruction Loss across model architectures. This heatmap shows the reconstruction loss (measured as the Mean Squared Error, MSE) between the real and generated data during model training and testing. The values correspond to the average MSE for the final 20 epochs of testing and are detailed in Sup. Fig. S1. The Y-axis represents the hidden layer dimensions, while the X-axis represents the latent space dimensions. B-D) Variational Autoencoder (VAE) model losses for the optimal architecture during training and testing. B) KL divergence loss. C) Reconstruction loss. D) Total loss. The blue lines represent the training loss, while the orange lines represent the test loss.

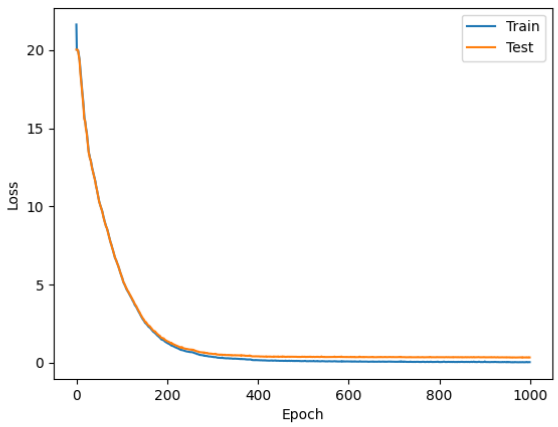


**Supplementary Figure S6. Reconstruction network losses.** The orange and blue lines represent the train and test losses, respectively, of the postprocessing network.

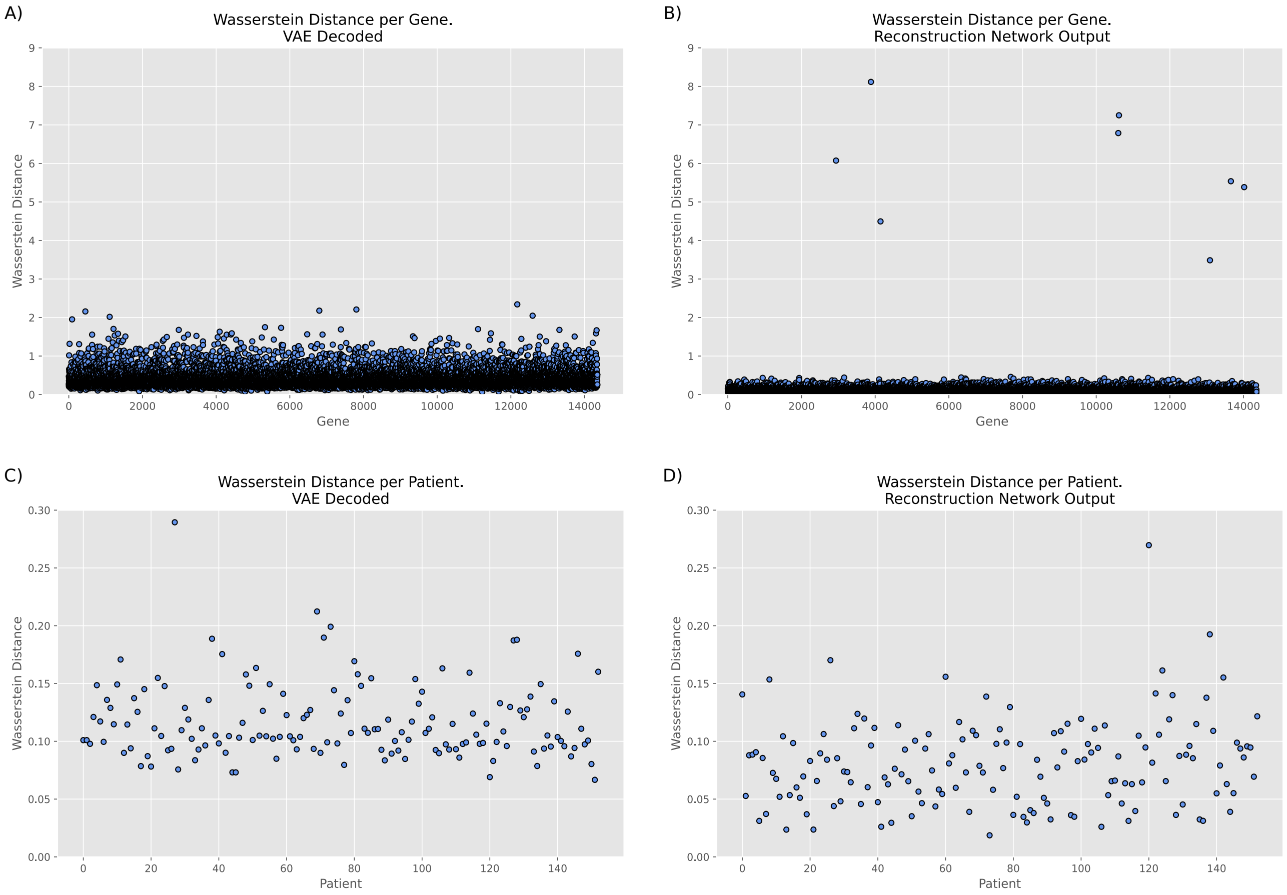


**Supplementary Figure S7. Reconstruction network performances.** Compared to VAE decoder output, the reconstruction network output shows a lower Wasserstein distance between the real and the generated data. Wasserstein distance between: (A, B) Genes, and (C, D) Patients. As shown in B, only eight genes (E2F3, SPRYD7, GDF5, FANCF, CECR6, RN7SKP101, VSIG8, DNM1P34) have a substantially high reconstruction error. These genes are neither among the most important for the data synthesis nor among the differentially expressed genes in G3, G3-G4, G4 subgroups. Notably, these genes are associated to highly variable biological functions, namely, cell cycle regulation (E2F3), association with leukemia (SPRYD7), extracellular matrix organization (GDF5), immune system response (FANCF), RNA binding activity (VSIG8). Among them, there is even a candidate gene (CECR6) and pseudogenes (RN7SKP101, DNM1P34). Potentially, the inherent variability of these genes may explain the difficulty in reconstructing them.

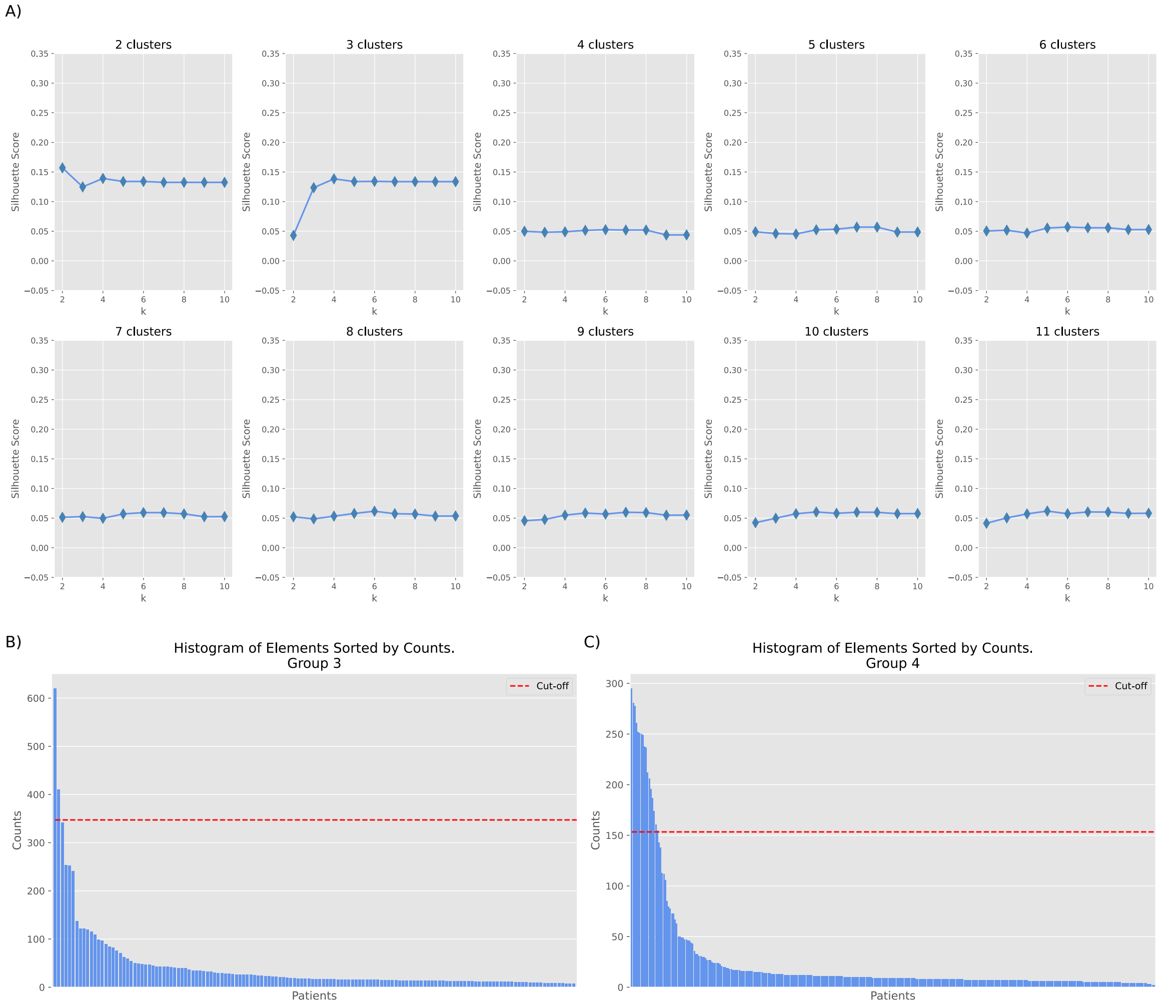


**Supplementary Figure S8. Identification of the G3-G4 patients. A) Silhouette scores for varying values of k neighbors and number of clusters.** This figure shows the silhouette scores for increasing numbers of k neighbors using k-Nearest Neighbors Graph (k-NNG) over increasing numbers of clusters obtained through agglomerative clustering. The preprocessed data was analyzed, focusing on Groups 3 and 4. Silhouette scores provide a measure of clustering consistency, with higher values indicating samples are better matched to their cluster. The results reflect the relationship between the number of neighbors k, the number of clusters, and the clustering appropriateness for the dataset. **B), C) Distribution of misclassified patients.** The histograms show the distribution of the number of times a patient was assigned to a cluster different from their true group, for B) Group 3 and C) Group 4. The red, dashed line represents a predefined cutoff; the expected value that randomly sampled patients would change subgroup (Methods). We simulate a coin-toss for the probability of a patient changing group. Data used is obtained after the preprocessing step.

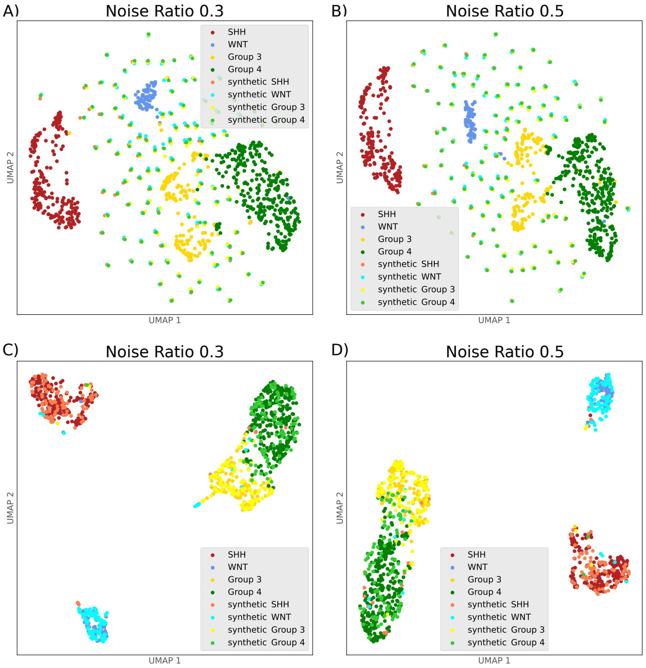


**Supplementary Figure S9. UMAPs comparing an alternative method for data generation with our VAE-assisted method.** A,B) show the UMAPs of the real data augmented with Gaussian noise. C,D) show the UMAPs of the latent space augmented with noise, then reconstructed. The sub-panels are organized by columns, representing increasing levels of added noise.

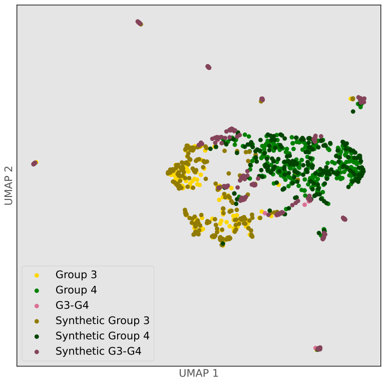


**Supplementary Figure S10. UMAP of augmented data.** The plot includes all real patients, and, overlaid, the synthetic ones in darker colors. The four original medulloblastoma subgroups, plus the newly-found G3-G4' subgroup, in lighter colors. Darker colors represent the synthetic patients. The noise ratio used for data generation was set to 0.2

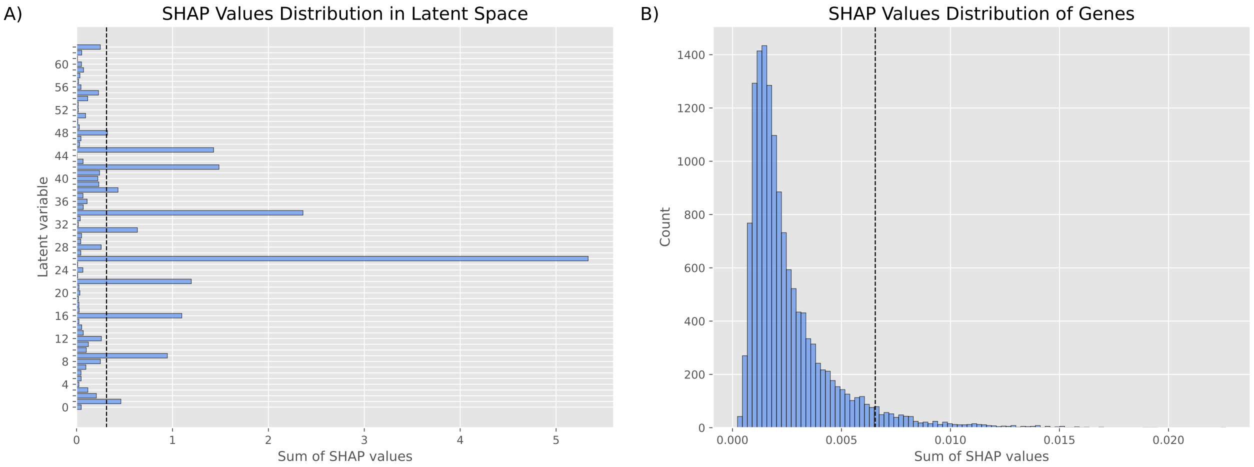


**Supplementary Figure S11. SHAP values distribution for the original medulloblastoma subgroup classification (SHH, WNT, G3, G4) in the latent space.** Distribution of the sum of SHAP values for A) the latent variables and B) the genes identified as the most important ones in the SHAP pipeline. The dashed, vertical line represents the cutoffs, defined in A) as the average of the sum of the SHAP values for the latent variables, and in B) as the top 5% genes with the highest SHAP values, i.e., those that explain most of the classification of the medulloblastoma subgroups. The most explainable latent variables from A), are used to obtain the genes in B) that better explain the subgroup classification.

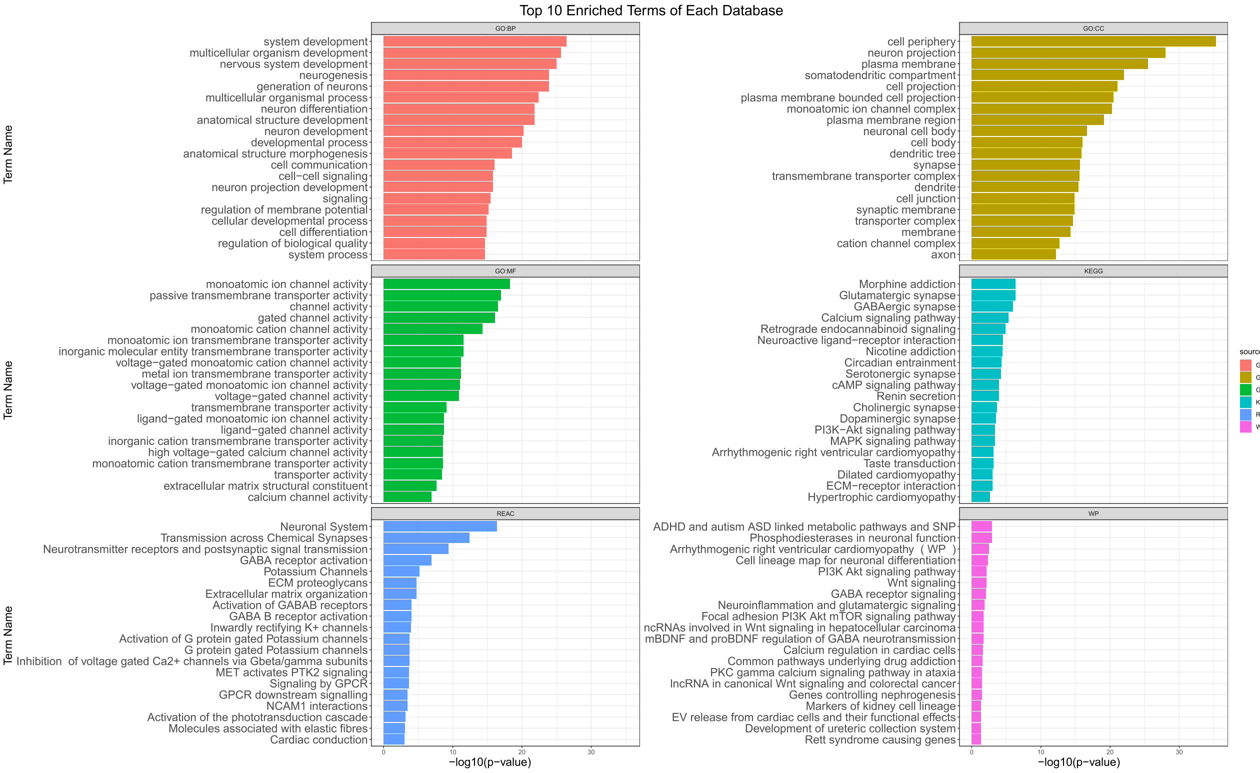


**Supplementary Figure S12. Enrichment analysis results on the top 5% most important genes from SHAP.** The figure shows the top 10 most significant terms and pathways for each of the databases considered. Each plot is titled with the name of the corresponding database: Gene Ontology (GO) (7), organized in its three aspects, Molecular Function (GO:MF), Cellular Component (GO:CC), and Biological Process (GO:BP); Reactome (REAC) (8); Kyoto Encyclopedia of Genes and Genomes (KEGG) (9); and WikiPathways (WP) (10). The enrichment analysis was conducted using *gprofiler* (4) on the genes identified as important (top 5%) for the classification of the original medulloblastoma subgroups in the VAE latent space (see Methods).

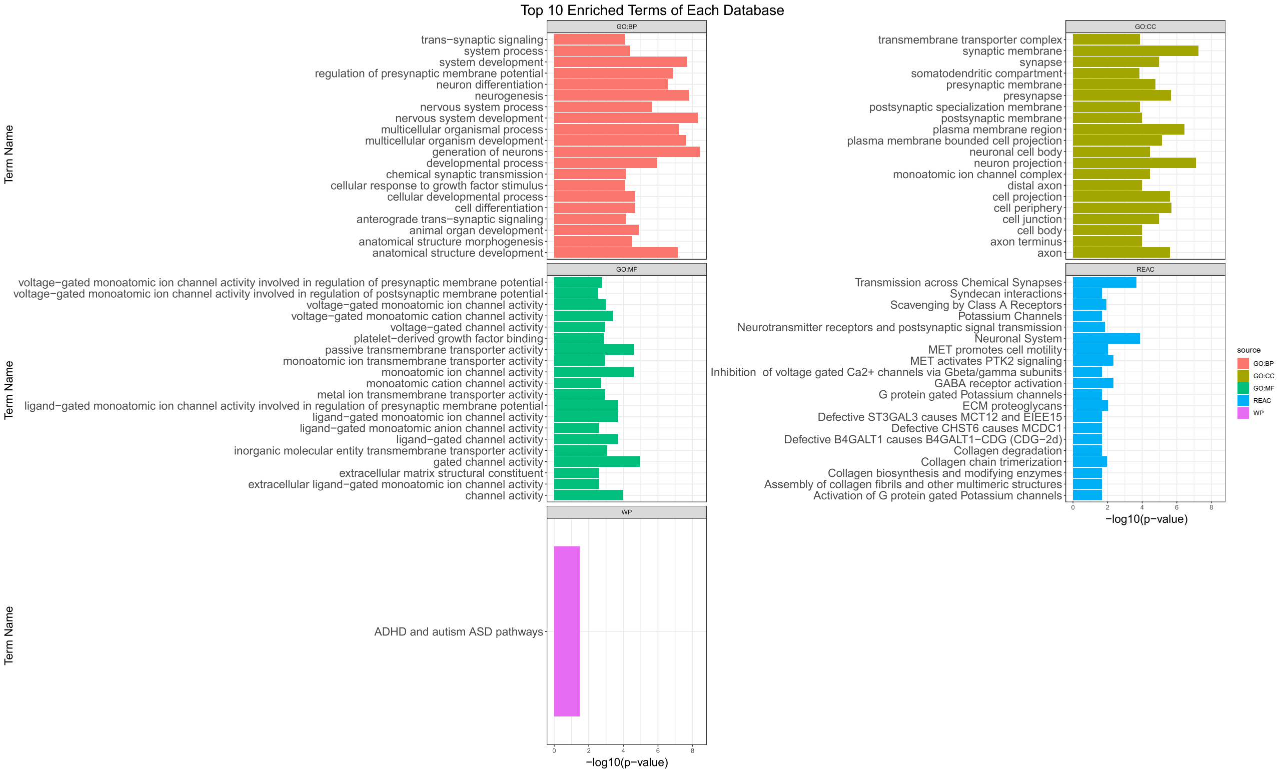


**Supplementary Figure S13. Enrichment analysis results on the top 1% most important genes from SHAP.** The figure shows the top 10 most significant terms and pathways for each of the databases considered, when 10 terms are available. Each plot is titled with the name of the corresponding database: Gene Ontology (GO) (7), organized in its three aspects, Molecular Function (GO:MF), Cellular Component (GO:CC), and Biological Process (GO:BP); Reactome (REAC) (8); and WikiPathways (WP) (10). Contrary to the previous analysis, the Kyoto Encyclopedia of Genes and Genomes (KEGG) (9) database did not return any terms. The enrichment analysis was conducted using *gprofiler* (4) on the genes identified as important (top 1%) for the classification of the original medulloblastoma subgroups in the VAE latent space (see Methods).

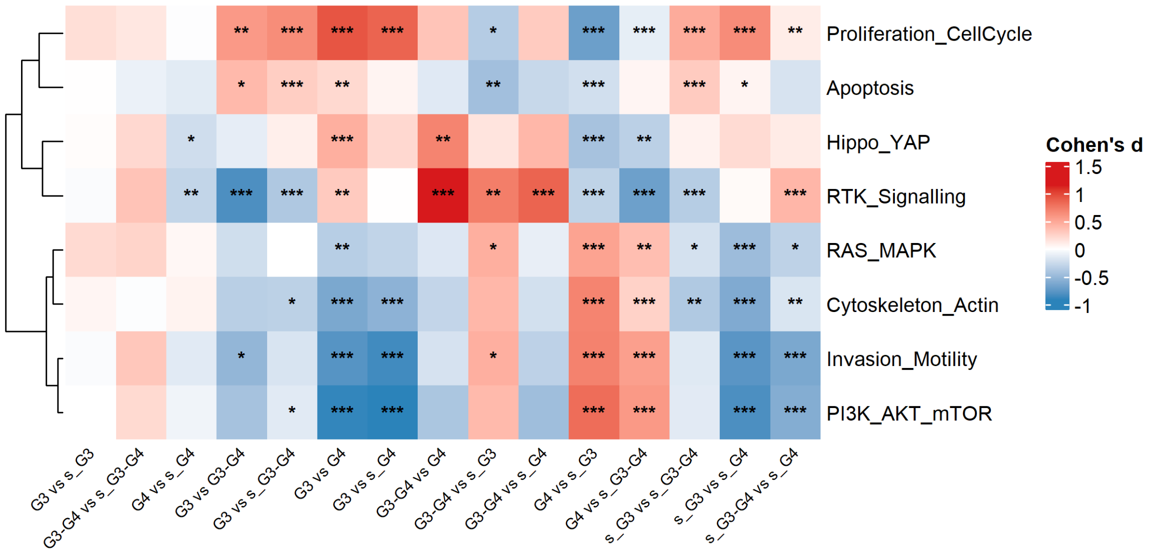


**Supplementary Figure S14. Heatmap of pairwise Cohen's d effect sizes across pathways and medulloblastoma groups.** Rows represent pathway activation scores; columns represent all pairwise comparisons between group classes. Cell colour encodes the signed Cohen's $d$ value using a diverging blue-white-red scale centered at zero, where positive values indicate higher pathway activation in Group1 relative to Group2, and negative values indicate the opposite. Asterisks within cells denote significance levels ($*p < 0.05$, $**p < 0.01$, $***p < 0.001$, Mann-Whitney-Wilcoxon test).

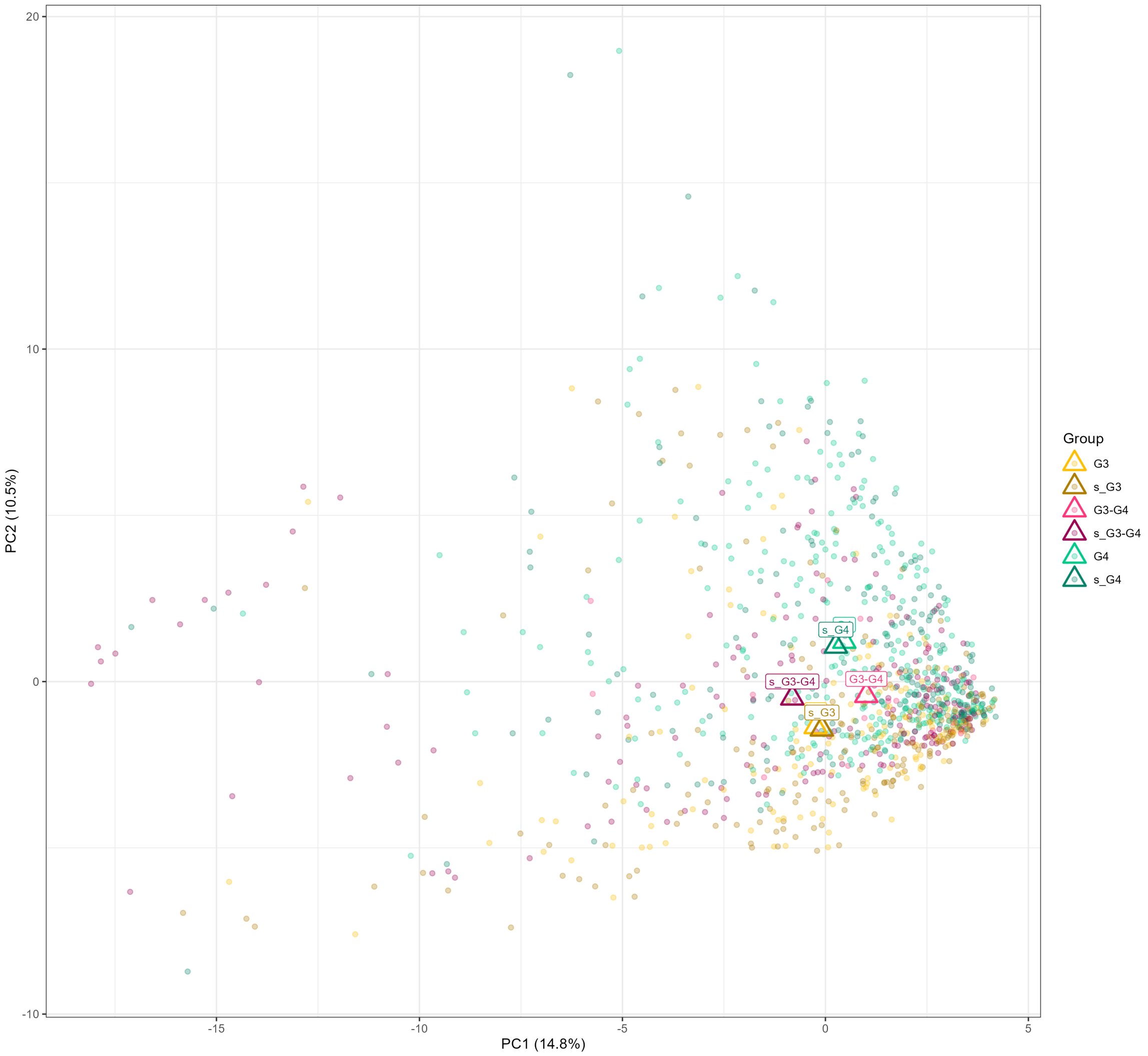


**Supplementary Figure S15. PCA of personalised simulations with subgroup and synthetic centroids.** PC1 and PC2 scores of all 1,363 personalised models are displayed, coloured by subgroup or corresponding synthetic class, with triangle symbols marking the centroid of each group. Centroids for G3 versus s_G3 and G4 versus s_G4 largely overlap, whereas G3–G4 and s_G3–G4 show a noticeable displacement along PC1 despite similar positions on PC2.

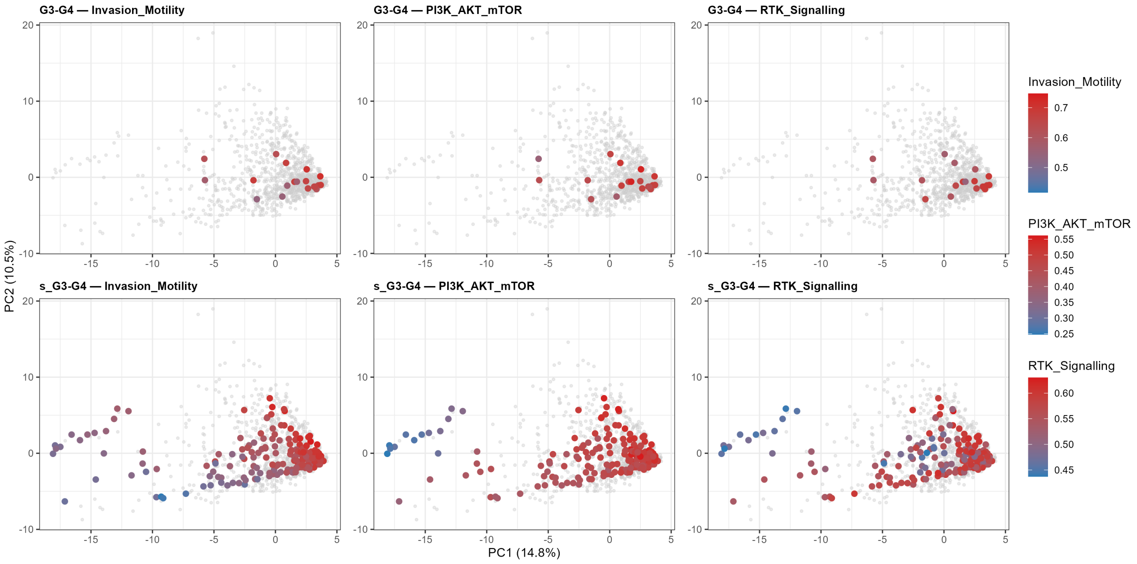


**Supplementary Figure S16. PCA of real and synthetic Group 3 and 4 models coloured by RTK, PI3K–AKT–mTOR and invasion/motility signalling.** PC1 and PC2 scores of all personalised models are shown, highlighting G3–G4 (top row) and s_G3–G4 (bottom row) samples, which are coloured according to invasion/motility, PI3K–AKT–mTOR or RTK signalling activity while other samples are shown in grey. Gradients along PC1 indicate that real and synthetic intermediate models occupy partially segregated regions that align with variation in these three pathways.

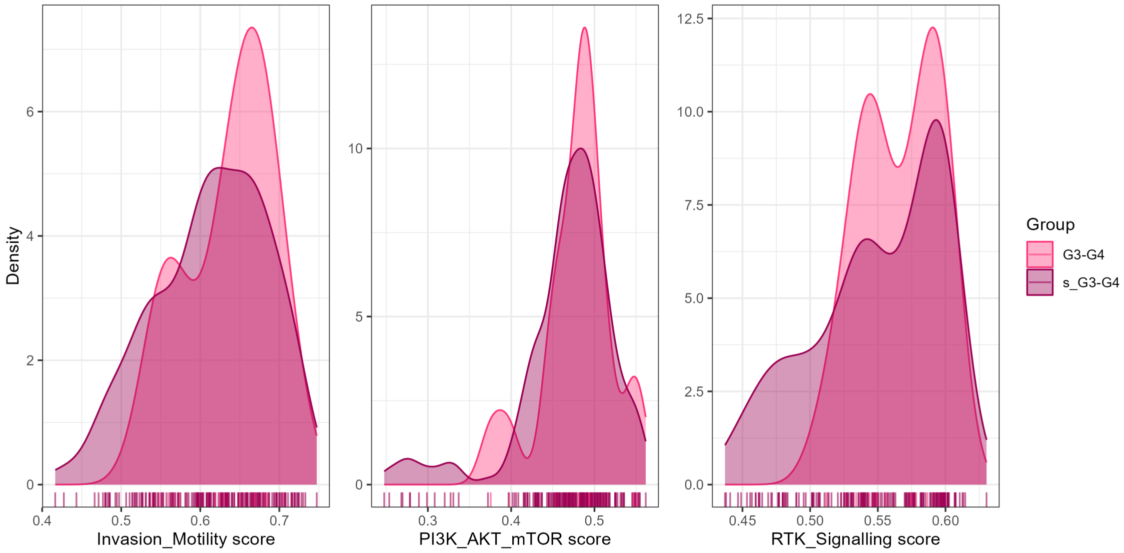


**Supplementary Figure S17. Distribution of key pathway scores in real versus synthetic G3–G4 models.** Densities of invasion/motility, PI3K–AKT–mTOR and RTK signalling scores are shown for G3–G4 and s_G3–G4 models, with rug plots indicating scores from individual patients' simulations. The two groups display broadly similar central tendencies but modest differences in dispersion and shape, particularly for RTK signalling
