## Supplementary figures and images for "Explainable Generative AI Uncovers a Molecular Continuum in Medulloblastoma with Implications for Rare Cancer Subtyping and Treatment Equity"

### MB_XARXA.png

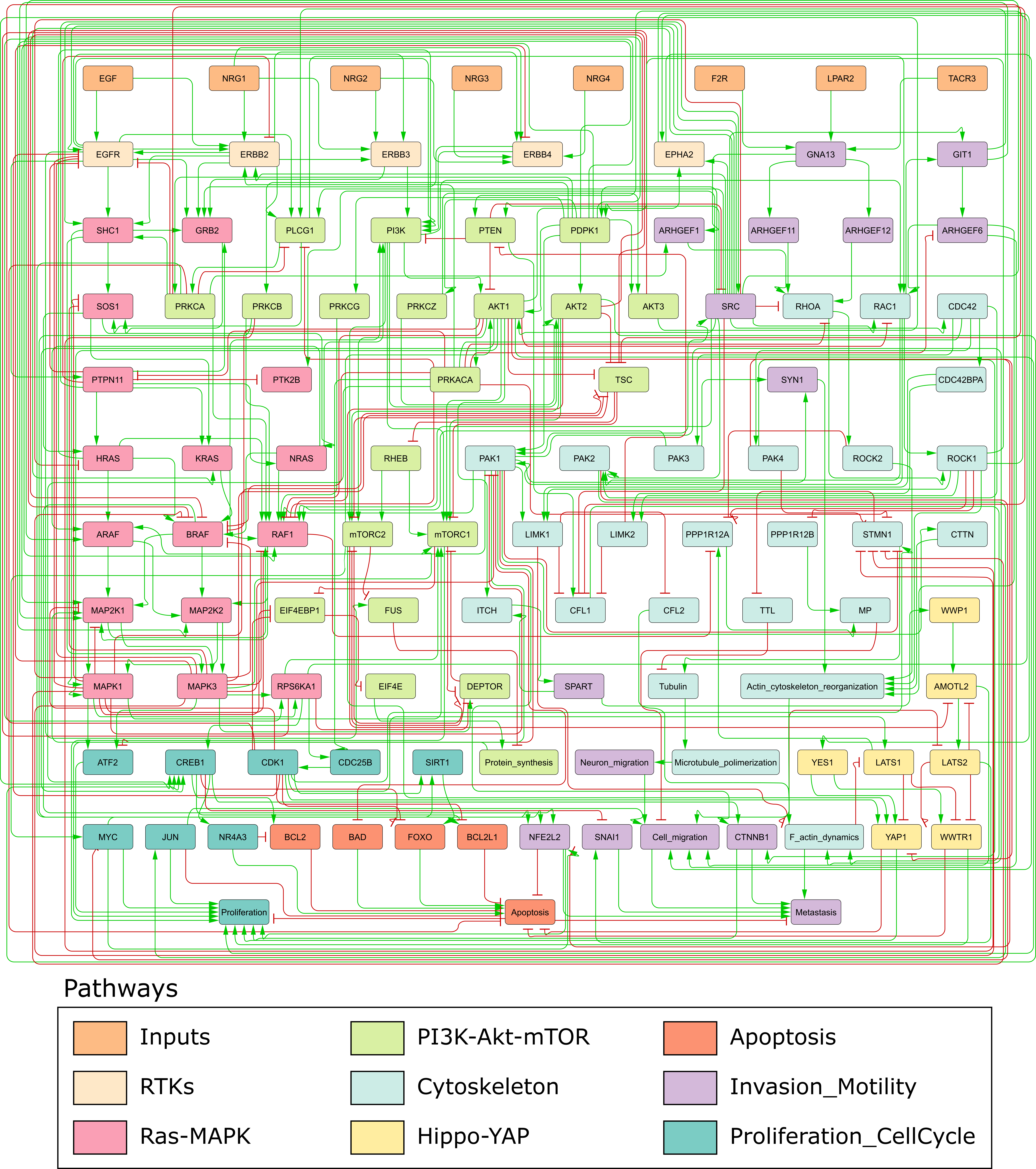

### MB_XARXA_blanc.png

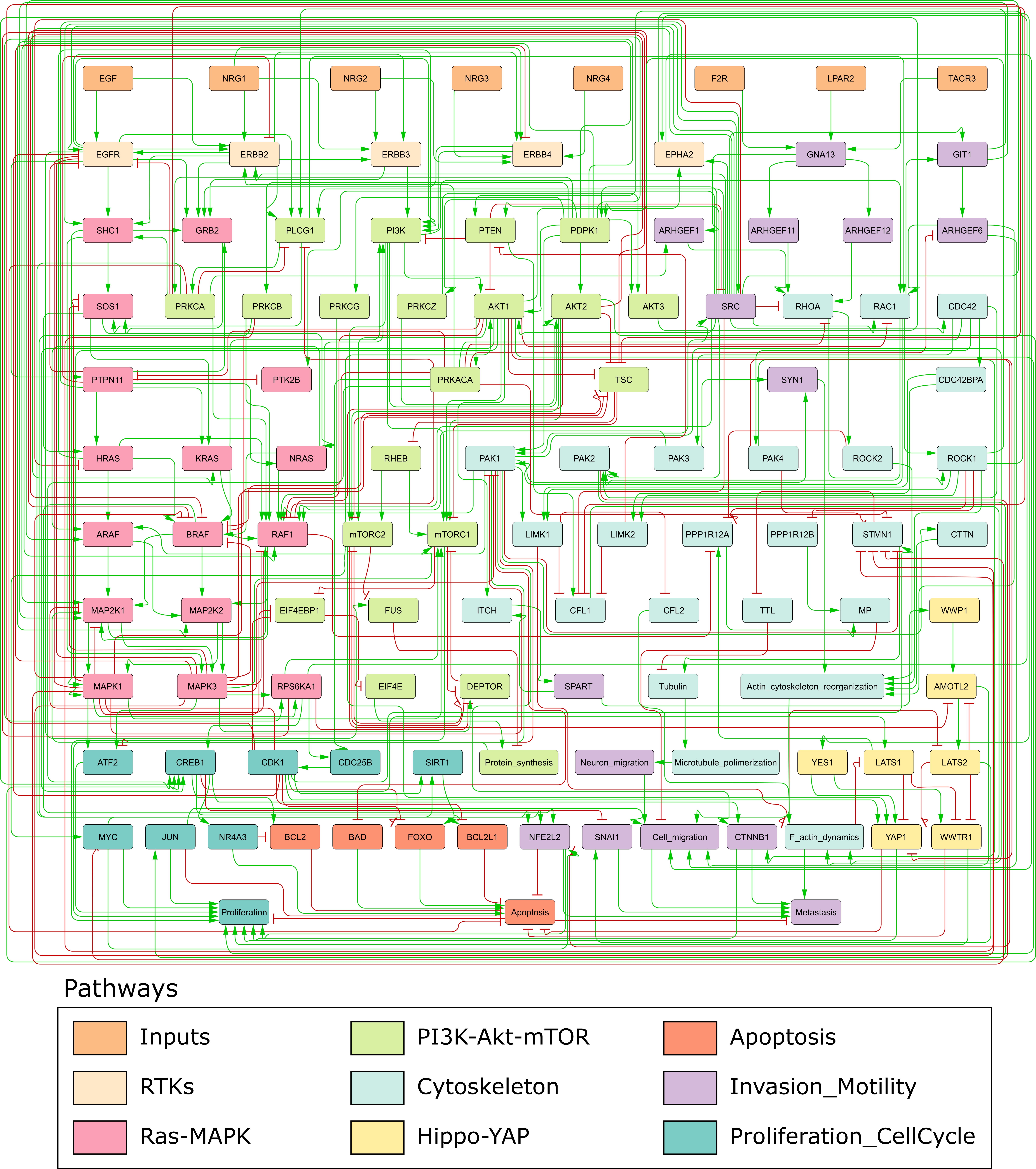
